# DNA methylation governs the sensitivity of repeats to restriction by the HUSH-MORC2 corepressor

**DOI:** 10.1101/2023.06.21.545516

**Authors:** Ninoslav Pandiloski, Vivien Horvath, Ofelia E. Karlsson, Georgia Christoforidou, Fereshteh Dorazehi, Symela Koutounidou, Jon Matas, Patricia Gerdes, Raquel Garza, Marie E. Jönsson, Anita Adami, Diahann Atacho, Jenny G. Johansson, Elisabet Englund, Zaal Kokaia, Johan Jakobsson, Christopher H. Douse

## Abstract

The human silencing hub (HUSH) complex binds to transcripts of LINE-1 retrotransposons (L1s) and other genomic repeats, recruiting MORC2 and other effectors to remodel chromatin. However, how HUSH and MORC2 operate alongside DNA methylation, a central epigenetic regulator of repeat transcription, remains poorly understood. Here we interrogate this relationship in human neural progenitor cells (hNPCs), a somatic model of brain development that tolerates removal of DNA methyltransferase DNMT1. Upon loss of MORC2 or HUSH subunit TASOR in hNPCs, L1s remain silenced by robust promoter methylation. However, genome demethylation and activation of evolutionarily-young L1s attracts MORC2 binding. Simultaneous depletion of DNMT1 and MORC2 causes massive accumulation of L1 transcripts. We identify the same mechanistic hierarchy at pericentromeric α-satellites and clustered protocadherin genes, repetitive elements important for chromosome structure and neurodevelopment respectively. Our data delineate the independent epigenetic control of repeats in somatic cells, with implications for understanding the vital functions of HUSH-MORC2 in hypomethylated contexts throughout human development.

## INTRODUCTION

Genetic elements may be repetitive either in the sense that there are many copies of the DNA sequence dispersed in the genome, the nucleotide sequence is itself repetitive, or both. Repeats are dynamic stretches of DNA, some of which can mobilize or duplicate. Taken together, repeats comprise more than half of the human genome.^1^ The most abundant are interspersed mobile elements called LINE-1 retrotransposons (L1s) that account for 17% of human DNA, a small subset of which continue to generate polymorphisms in the population via retrotransposition events.^2, 3^ Non-coding, low-complexity repeats such as alpha-satellites (ALRs), which occupy up to 5% of the genome, are not mobile but like L1s are hotspots of recombination and drive genomic complexity within and between individuals.^3–5^

Aside from being sources of genetic variation, the functional importance of repeats is increasingly recognized as technologies to manipulate and resolve their sequences improve. For example, even though the vast majority are retrotransposition-incompetent,^3^ L1s impact transcriptional programs by acting as alternative promoters and via changes in chromatin structure^6^ – thereby rewiring gene expression^7^ and contributing to functional diversification in tissues including the human brain.^8^ ALRs form higher-order structures at centromeres that are critical to the structural integrity of chromosomes, while other tandem repeats have widespread effects on gene transcription.^9, 10^ It is also notable that many protein and RNA genes are themselves arrayed in clusters, where the genetic structure may be functionally significant.^11^ For example, the repetitive array of exons in clustered protocadherin genes are expressed combinatorially, generating a barcoding system that underpins neuronal individuality and complexity in the nervous system.^12^

The activity of repeats and repetitive genes is tightly regulated since aberrant control of these elements is associated with genomic instability and disease.^13–16^ Packaging of repeats into particular chromatin states is a conserved way to control their transcription and replication. In somatic human cells, repeats are decorated by patterns of DNA and histone methylation that are established early in development.^17, 18^ Promoter CpG methylation, maintained in somatic cells by DNA methyltransferase 1 (DNMT1) – and trimethylation of lysine 9 in histone H3 (H3K9me3), maintained by histone methyltransferases such as SETDB1, are enriched over repeats and correlate with heterochromatin formation and transcriptional repression.^19, 20^

The human silencing hub (HUSH) complex^21^ has emerged as an important epigenetic regulator of repeats through H3K9me3.^7, 22–24^ Composed of TASOR, MPP8 and Periphilin, HUSH recruits the nuclear ATPase MORC2 and H3K9me3 writer SETDB1 to remodel chromatin.^21, 25^ Recent data have shown that HUSH also participates as an adapter for co-transcriptional RNA processing complexes, leading to the destruction or termination of targeted transcripts.^26–28^ Such an RNA-dependent mechanism is reminiscent of evolutionarily-ancient systems such as the yeast RNA-induced transcriptional silencing (RITS) complex, and indeed analysis of HUSH protein architectures revealed striking similarities with RITS.^22^ Endogenous genomic repeats targeted by the HUSH-MORC2 co-repressor – that is, L1s and certain repetitive genes^7, 22, 25, 29^ – appear to be unified by the presence of long, intronless transcriptional units.^30^ Experiments with transgene reporters have shown that transcription of such units is required for recruitment of the complex.^7, 27, 30^ Nonetheless, how the HUSH-MORC2 axis interacts with DNA methylation, the central epigenetic regulator of endogenous repeat transcription in somatic cells, is poorly understood. Key mechanistic studies of HUSH-MORC2 have been done in cancer lines that are characterized by aberrant DNA methylation – or mouse embryonic stem cells, which are hypomethylated (depending on culture conditions^31^) and whose repetitive genome diverges substantially from humans.^32^ Furthermore, since deletion of DNA methyltransferase 1 (DNMT1), the enzyme that maintains CpG methylation during cell division, is lethal in cancer cells and somatic mammalian cells,^33–35^ experiments that provide mechanistic insights into the relationship between HUSH-MORC2 and DNA methylation in repeat regulation are challenging to design.

Here we test the relationship between HUSH-MORC2 and DNA methylation in embryo-derived human neural progenitor cells (hNPCs), a somatic model of brain development with robust levels of DNA methylation that nonetheless tolerates acute removal of DNMT1 and global genome demethylation.^36^ Loss of MORC2 or defining HUSH subunit TASOR in hNPCs does not cause widespread misexpression of L1s, all but a handful of which are maintained in a transcriptionally silent state by promoter DNA methylation. However, upon removal of DNA methylation and resulting transcriptional activation, these elements accumulate MORC2 and are sensitized to HUSH-MORC2 restriction such that simultaneous depletion of DNMT1 and MORC2 leads to massive accumulation of LINE-1 transcripts. We provide evidence of the same mechanistic hierarchy at pericentromeric ALRs and clustered protocadherin genes, repetitive elements important for chromosome stability and nervous system development respectively. Together our data delineate the independent epigenetic control of repeats in somatic cells, with implications for understanding the vital functions of the HUSH-MORC2 axis in hypomethylated contexts throughout human development.

## RESULTS

### Epigenomic profiling of the human neural progenitor cell (hNPC) model

To investigate the relationship of the HUSH-MORC2 axis with DNA methylation in epigenetic repeat regulation (**Fig. 1A**) we used a somatic human neural progenitor cell (hNPC) line derived from fetal brain 6 weeks post-conception,^37^ which represents a timepoint of post-implantation embryonic development when DNA methylation has been fully re-established.^17, 36^ These cells have the characteristics of human neuroepithelial-like stem cells.^37^ We confirmed expression of the NPC marker Nestin and the neural differentiation capacity of the cells (**Fig. 1B, Supp. Fig. 1A**). Analysis of the hNPC proteome detected all core HUSH complex components (TASOR, MPP8, PPHLN1), its effectors (MORC2, SETDB1) and maintenance DNA methyltransferase DNMT1 (**Supp. Fig. 1B**).^36^ Single nuclei RNA-seq analysis^8^ of six fetal forebrain tissue samples (7.5-10.5-weeks post-conception) illustrated widespread expression of the genes encoding these factors in all identified cell types of the developing human brain (**Fig. 1C, Supp. Fig. 1C**).

**Figure 1.**
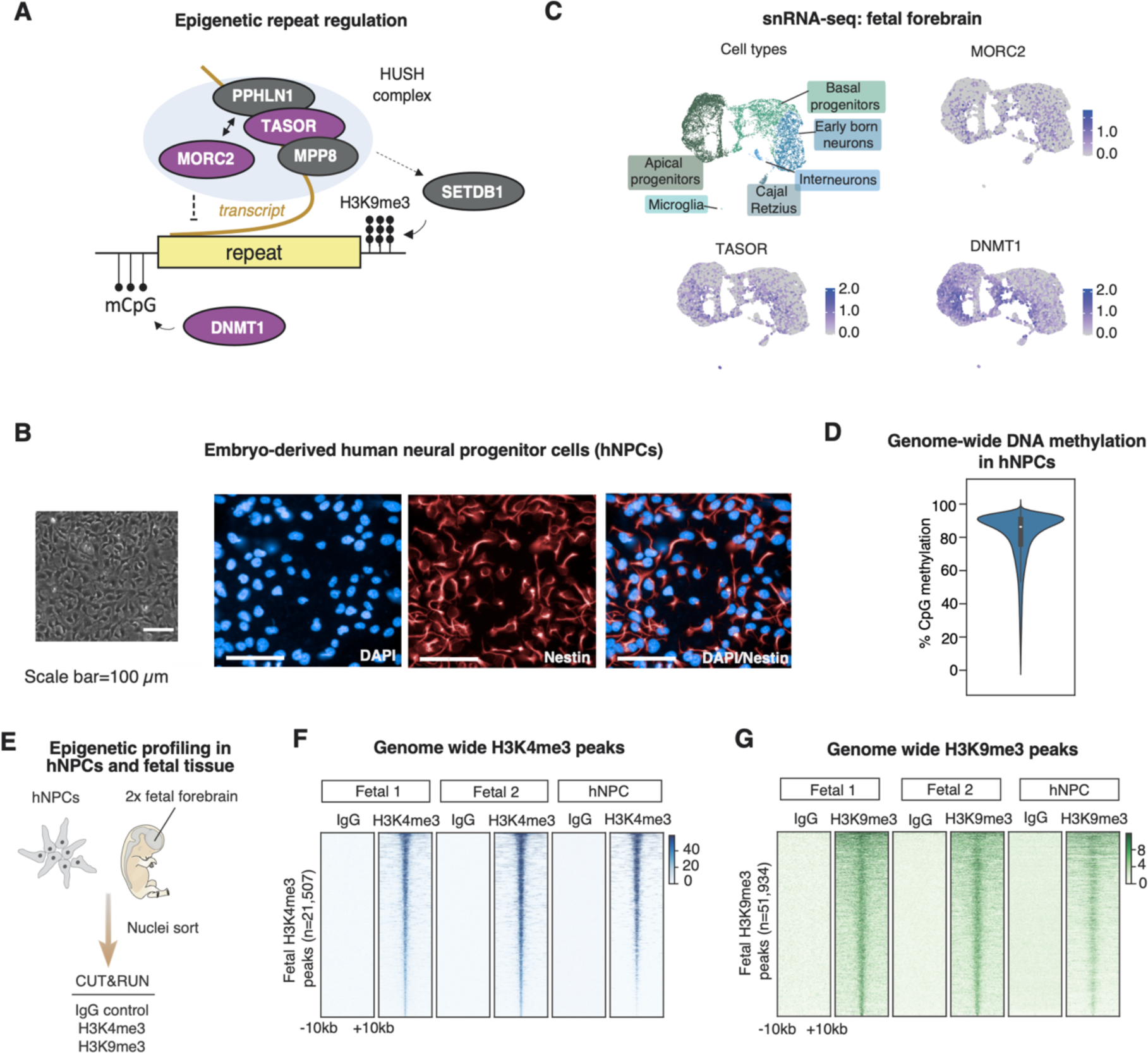
Epigenetic control of repeats in a somatic model of human brain development. **(A)** Simplified model of epigenetic transcriptional regulation of genomic repeats by DNA methylation and the HUSH-MORC2 corepressor. Factors shaded in purple are subject of particular focus in this study. (B) Brightfield image (left) and immunostaining (right) of the hNPC line used in this study. **(C)** Expression of MORC2, TASOR and DNMT1 in the developing human forebrain according to snRNA-seq analysis (n=6).^8^ (**D)** Assessment of genome-wide CpG methylation according to Oxford Nanopore whole-genome sequencing of the hNPC line in 10-kb bins. **(E**) Schematic of CUT&RUN epigenome profiling experiments in human fetal forebrain tissue (n=2) and hNPCs (n=3). **(F, G)** Heatmaps illustrating CUT&RUN signal enrichment of H3K4me3, H3K9me3 and a non-targeting IgG control in fetal forebrain samples and hNPCs plotted over consensus peaks called in fetal samples. Displayed are the genomic regions spanning +/- 10kb from the peak centre.

Given our focus on repetitive elements in this study we used long-read Oxford Nanopore sequencing to resolve DNA methylation patterns in the hNPC genome.^38^ We found that more than 80% of CpGs are methylated genome-wide (**Fig. 1D**). To investigate the histone methylation status of the hNPC model and compare it to actual human tissue, we performed CUT&RUN profiling of H3K4me3 (enriched at active transcription start sites) and heterochromatin mark H3K9me3 in hNPCs and human fetal forebrain samples (n=2, 7- and 10-weeks post conception) alongside non-targeting IgG controls (**Fig. 1E**). More than 90% of H3K4me3 and H3K9me3 peaks called in fetal tissue samples were also occupied by H3K4me3 and H3K9me3 in the hNPC model, and vice versa (**Fig. 1F,G, Supp. Fig. 1D,E**). Together these experiments suggest that hNPCs represent a good cellular model for studying epigenetic mechanisms of repeat regulation in human brain development and confirm that the DNA methylation patterns of somatic human cells are present in the hNPC model.

### Targeted depletion of the HUSH and MORC2 with CRISPRi

To inhibit expression of MORC2 or defining HUSH subunit TASOR^22^ we used a bulk lentiviral CRISPR interference (CRISPRi) approach. We co-expressed a dCas9-KRAB-GFP fusion transcript with gRNAs targeted to the relevant TSS, or a non-targeting control (**Fig. 2A**). Underpinning CRISPRi is chromatin remodelling of the targeted TSS through KRAB repressor binding, accumulation of promoter H3K9me3 and transcriptional silencing. Following 10 days’ expansion, epigenome profiling in the CRISPRi lines indeed revealed targeted accumulation of H3K9me3, with concomitant loss of H3K4me3 over the relevant TSS (**Fig. 2B**). This corresponded to essentially complete depletion of MORC2 and TASOR transcript (**Fig. 2C**) and protein (**Fig. 2D**) levels. Neither MORC2 nor TASOR depletion caused an obvious morphological change in hNPCs and Nestin staining was consistent across treatment and control groups, suggesting that cell identity was not perturbed by loss of these factors (**Fig. 2E**). Thus, CRISPRi proved a successful strategy for targeted and acute depletion of HUSH and MORC2 in hNPCs.

**Figure 2.**
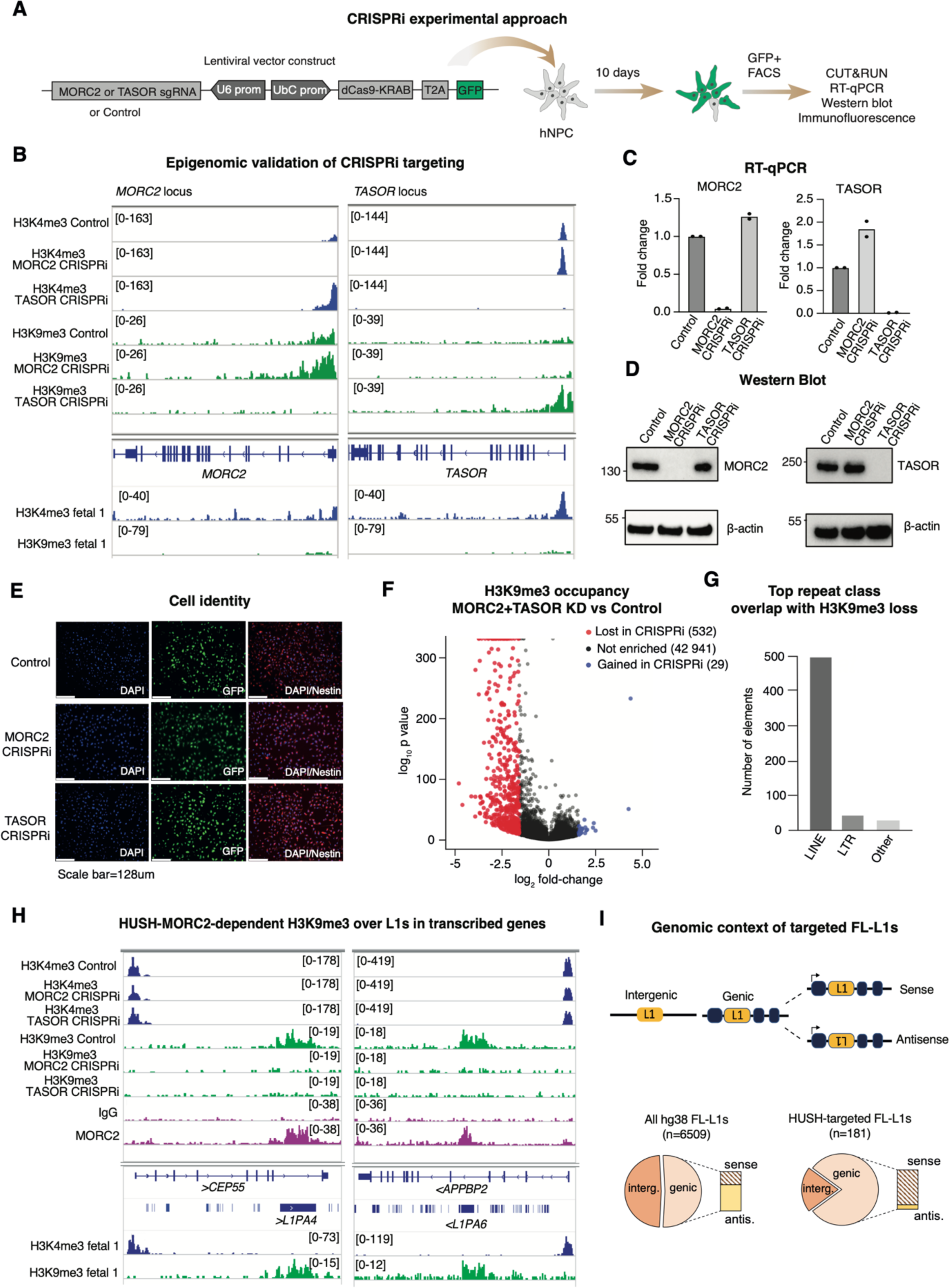
The HUSH-MORC2 pathway is active in hNPCs. **(A)** Schematic of CRISPRi method to deplete MORC2 or HUSH subunit TASOR. **(B)** Genome browser snapshots of *MORC2* (left) and *TASOR* (right) genes illustrating epigenome editing by the CRISPRi technology, as measured by CUT&RUN sequencing (H3K4me3, H3K9me3) relative to non-targeting control. Tracks from a fetal forebrain sample are shown for comparison. **(C,D)** Reverse transcription quantitative PCR (RT-qPCR) and Western blot analysis of MORC2 (left) and TASOR (right) transcript and protein levels in MORC2 and TASOR CRISPRi cells, relative to a non-targeting control, 10 days post-transduction. **(E)** Immunostaining to assess dCas9-KRAB-T2A-GFP and Nestin expression in MORC2 and TASOR CRISPRi cells. (**F)** Volcano plot showing H3K9me3 changes genome-wide over H3K9me3 peaks in MORC2 (n=2) and TASOR (n=2) CRISPRi hNPCs compared to controls (n=4) as measured by CUT&RUN epigenome profiling 10 days post-transduction (significant points defined by fold-change>3 and Poisson p-value <1e-04). **(G)** Top repeat class overlaps with regions losing H3K9me3 upon MORC2 and TASOR depletion. **(H)** Genome browser snapshot examples of intronic L1PA elements occupying the same strand as a transcribed host gene, bound by MORC2 and losing H3K9me3 upon MORC2 and TASOR depletion. **(I)** Summary of genomic attributes of FL-L1s losing H3K9me3 upon MORC2 and TASOR depletion.

Given the well-established association of the HUSH-MORC2 corepressor with H3K9me3 maintenance,^7, 21, 22^ we assessed changes in H3K9me3 genome-wide, pooling replicates from MORC2 and TASOR CRISPRi experiments. We observed loss of H3K9me3 over hundreds of sites, confirming the pathway as functional in the hNPC model (**Fig. 2F,G**). CUT&RUN using an optimized crosslinking protocol (**Supp. Fig. S2A**) showed MORC2 bound to the majority of these sites (**Fig. 2H**, **Supp. Fig. 2B**). Repeats marked by H3K9me3 in a HUSH-MORC2-dependent manner in hNPCs were predominantly L1s in transcribed genes (**Fig. 2H,I**). Notably, 88% of these occupied the same strand as the host gene, a striking observation given that intronic L1s predominantly lie in the opposite orientation to their host genes (outweighing sense insertions by about 2:1)^39^ (**Fig. 2I**). Targeting of sense insertions in transcribed genes supports the model that transcription of the A-rich L1 forward strand is a determinant of HUSH recognition and H3K9me3 deposition, regardless of whether transcription is element-derived (i.e. initiated at the L1 promoter, see below) or from an upstream gene promoter.

### DNA methylation, but not HUSH-MORC2, controls L1 transcription in hNPCs

Full-length L1s (FL-L1s, here defined as > 6 kb) contain a promoter in their 5’ UTR that drives production of a bicistronic mRNA encoding two open reading frames (ORF1p and ORF2p, **Fig. 3A**). Since L1 integrants accumulate mutations over evolutionary time, elements can be dated and classified according to evolutionary age, including human-specific (L1HS) and hominoid-specific (L1PA2-L1PA4) subfamilies. FL-L1s from these youngest subfamilies retain the capacity to be actively transcribed, but are efficiently silenced by promoter DNA methylation in hNPCs.^36^ Clones of human embryonic stem cells or immortalized cancer lines that lacked components of HUSH or MORC2 previously showed pronounced upregulation of young L1s, an interferon response and production of L1-derived protein ORF1p.^7, 22, 29, 40^ However, ESCs and cancer cells are characterized by incomplete or aberrant DNA methylation. To investigate transcriptional regulation of L1s by HUSH-MORC2 in hNPCs, where L1s are robustly methylated (**Fig. 3B**), we used a 2×150bp, polyA-enriched stranded library preparation for bulk RNA-seq following MORC2/TASOR CRISPRi, with a reduced fragmentation step to optimize library insert size for repeat analysis. Using this approach reads can be mapped uniquely to most individual L1 instances, except for a few of the youngest L1s and polymorphic alleles that are not annotated in the reference assembly.^8, 36^ In differential expression analysis of L1s and other repeats we routinely used two approaches – ‘unique mapping’, where ambiguously mapped reads were discarded, and ‘multi-mapping’ where those reads were retained and assigned to subfamilies by the TEtranscripts software.^41^ As a control we took advantage of a validated CRISPR vector to disrupt the catalytic domain of DNMT1, known to activate evolutionarily-young FL-L1s (**Fig. 3C**).^36^

**Figure 3.**
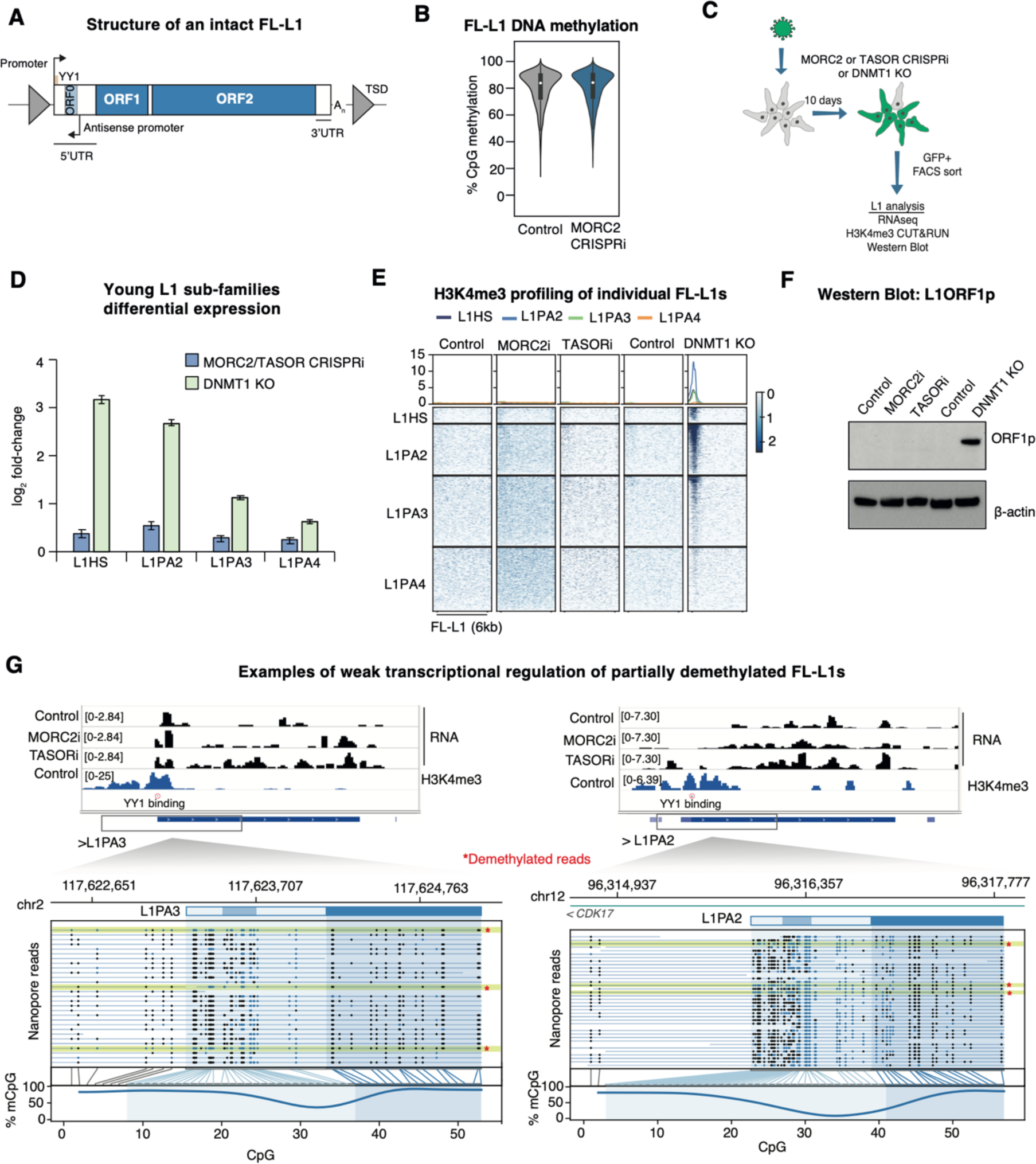
FL-L1s are generally silenced by DNA methylation but not HUSH-MORC2 in hNPCs. **(A)** Genetic structure of a full-length (>6-kb) L1 retrotransposon. TSD, target site duplication **(B)** Nanopore DNA methylation analysis over reference FL-L1s in control (n=1) and MORC2 CRISPRi hNPCs (n=1). (B) Schematic of L1 analysis of MORC2 and TASOR CRISPRi and DNMT1-KO hNPCs. **(D)** Average log_2_-fold-change (LFC) of young L1 subfamilies measured by RNA-seq in MORC2 or TASOR CRISPRi hNPCs (n=7) versus controls (n=4) using the TEtranscripts software. DNMT1-KO RNA-seq data in hNPCs taken from Jönsson et al.^36^ In each case the error bars represent +/- standard error of the mean calculated by DEseq2. **(E)** Heatmaps illustrating CUT&RUN signal enrichment of H3K4me3 in control, MORC2 CRISPRi, TASOR CRISPRi and DNMT1-KO hNPCs, plotted over young full-length L1PA families sorted by evolutionary age (top to bottom). Only signal with mapQ score > 10 was used to generate signal matrices. In all cases experiments were performed 10 days post-transduction. **(F)** Western blotting for L1 orf1p expression in control, MORC2 CRISPRi, TASOR CRISPRi and DNMT1-KO hNPCs. **(G)** Two examples of FL-L1s weakly upregulated by MORC2 and TASOR CRISPRi showing promoter H3K4me3 in control cells and an intact YY1 binding site (upper). Nanopore sequencing reads suggest incomplete DNA methylation over the promoter in control hNPCs (lower).

Strikingly, the widespread loss of H3K9me3 we observed over L1s was not accompanied by widespread transcriptional deregulation of L1s in hNPCs lacking MORC2 or TASOR: we detected very modest transcriptional changes amongst young L1 subfamilies (including multimapped reads) and in genes harbouring HUSH-MORC2-targeted FL-L1PAs (**Fig. 3D, Supp. Fig. 3A**). CpG methylation status over L1s in cells lacking MORC2 remained unchanged (**Fig. 3B**). In accordance with a lack of transcriptional changes, we did not detect accumulation of H3K4me3 over the 5’ UTR promoter of young FL-L1s upon TASOR or MORC2 depletion (**Fig. 3E**), nor an interferon response (**Supp. Fig. 3B**). We did not detect ORF1p expression in TASOR or MORC2 CRISPRi cell lines by Western blotting (**Fig. 3F**). By contrast, following disruption of DNMT1,^36^ pronounced transcriptional activation of FL-L1s was accompanied by H3K4me3 over FL-L1 promoters and ORF1p expression (**Fig. 3D-F**). Notably, H3K9me3 was not significantly affected over young FL-L1s in the DNMT1-KO cells over the 10-day timeframe of the experiment (**Supp. Fig. 3C**), suggesting that H3K9me3 remains despite expression of these elements.

In accordance with subfamily-wide results, when unique mapping was enforced in transcriptomic analysis, we found that only 19 individual reference FL-L1s were significantly upregulated upon MORC2 or TASOR CRISPRi (**Supp. Fig. 3D**). Analysis of the sequences and epigenetic status of these elements showed that only three had both an intact YY1 binding site (required for accurate transcription initiation^42^) and evidence of H3K4me3 enrichment over their promoter, suggesting that the remaining elements were transcribed from other promoters lying upstream of the L1. Interestingly, upon examination of the promoters of the three candidate elements in our Nanopore sequencing dataset we could detect a proportion of demethylated reads (**Fig 3G** and **Supp. Fig. 3E**), suggesting that these elements are incompletely methylated in hNPCs.

Together our results illustrate that loss of HUSH-MORC2 is not sufficient to activate L1 transcription in hNPCs, which is silenced by promoter DNA methylation. In the small handful of cases where DNA methylation over the L1 promoter is incomplete in hNPCs, this correlates with leaky transcriptional activity at steady-state and evidence of HUSH-MORC2-dependent transcriptional regulation at those loci.

### DNA methylation status controls L1 sensitivity to HUSH-MORC2 restriction

We therefore hypothesized that DNA methylation status, by controlling L1 promoter activity, determines the sensitivity of FL-L1s to HUSH-MORC2 restriction. To test this, we first assessed MORC2 binding to L1s upon genome demethylation (**Fig. 4A**). We used CRISPRi as an alternative approach to DNMT1 knockout reported previously, validating two gRNAs for successful inhibition of DNMT1 transcription by RT-qPCR (**Supp. Fig 4A**). Global loss of DNA methylation in DNMT1 CRISPRi cells was confirmed by 5mC immunostaining (**Supp. Fig. 4B**). RNA-seq analysis showed pronounced derepression of the evolutionarily-youngest L1PA families and imprinted or germline-restricted genes, consistent with DNMT1-KO hNPCs (**Supp. Fig 4C,D**).^36^ Upon depletion of DNMT1 we could measure clear accumulation of MORC2 over activated, young FL-L1s (**Fig. 4A**). Next, to test the functional impact of MORC2 on the transcription of demethylated L1s, we designed double CRISPRi experiments to deplete DNMT1 in combination with MORC2 (**Fig. 4B**). We compared the transcriptomes of the MORC2/DNMT1 double CRISPRi treatment to non-targeting controls and these datasets to individual MORC2 and DNMT1 CRISPRi experiments. Loss of MORC2 and DNMT1 expression was confirmed in the RNA-seq data (**Fig. 4C**). Using unique mapping to discriminate FL-L1s from truncated copies, we observed upregulation of 2,504 elements (LFC >2, padj <0.05) upon simultaneous loss of MORC2 and DNMT1, more than a third of all FL-L1s in the hg38 reference genome. The expression of L1s from human- and primate-specific subfamiles (L1HS, L1PA2, L1PA3, L1PA4) was dramatically increased in the double CRISPRi experiment, whether the analysis was done at a subfamily level (**Fig. 4D**) or at an individual element level (considering only FL-L1s and using unique mapping) (**Fig. 4E,F**). Importantly, the deregulation of long terminal repeat (LTR) transposon families (e.g. LTR12C, Harlequin) and imprinted or germline-restricted genes (e.g. *XIST, DAZL, MAEL, NNAT*) – activated in cells lacking DNMT1 but not thought to be controlled by HUSH-MORC2 – were unchanged between the DNMT1 and MORC2/DNMT1 CRISPRi treatments (**Supp. Fig. 4E,F**). We did though observe evidence of compounded effects of the double CRISPRi at several L1-fusion genes^36^ i.e. where a young L1 provides an alternative promoter for a gene (e.g. *GJB7, GABRR1, WDR72*) (**Supp. Fig. 4G**).

**Figure 4.**
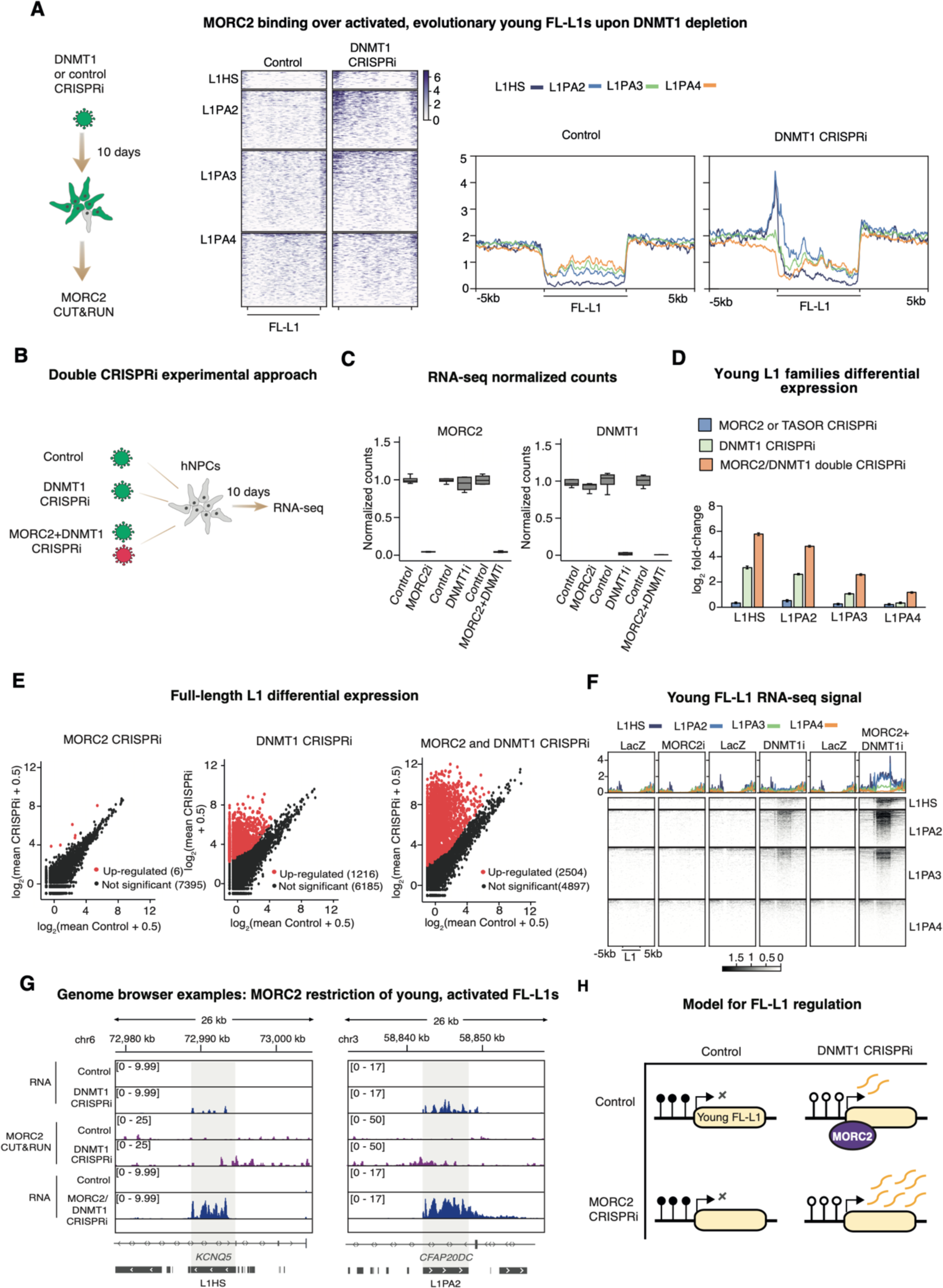
Genome demethylation sensitizes young L1s to restriction by MORC2. **(A)** Measurement of MORC2 L1 binding upon genome demethylation by DNMT1 CRISPRi. Heatmaps illustrating CUT&RUN signal enrichment of MORC2 and IgG controls in control and DNMT1 CRISPRi hNPCs, plotted over young full-length L1 subfamilies sorted by evolutionary age (top to bottom). Summary plots are shown (right). Only uniquely-mapped reads were used to generate signal matrices. Displayed are the genomic regions spanning +/- 5kb from the peak centre. **(B)** Strategy for simultaneous MORC2 and DNMT1 CRISPRi. **(C)** Box plots showing normalized read counts of bulk RNA-seq experiments in control (n=4), MORC2 CRISPRi (n=3), control’ (n=4), DNMT1 CRISPRi (n=4), control’’ (n=4) and MORC2/DNMT1 double CRISPRi (n=4) hNPCs **(D)** Average log_2_-fold-change (LFC) of young L1 subfamilies measured by RNA-seq in MORC2 or TASOR CRISPRi hNPCs (n=7, blue), DNMT1 CRISPRi (n=4, green) and MORC2/DNMT1 double CRISPRi (n=4, orange) versus controls (n=4 for each independent experiment) using TEtranscripts. In each case the error bars represent +/-standard error of the mean calculated by DEseq2. **(E)** Mean plots of RNA-seq read counts for different treatments versus their respective controls over reference FL-L1s. Red points indicate that the element satisfied both log_2_-fold-change >2 and padj < 0.05 cutoffs as calculated by DEseq2 where unique mapping was enforced. **(F)** Heatmaps of RNA-seq signal plotted over young full-length L1PA families sorted by evolutionary age (top to bottom). **(G)** Examples of young FL-L1 transcriptional changes upon DNMT1 and MORC2/DNMT1 depletion. MORC2 binding in DNMT1 CRISPRi cells is shown. **(H)** Model for independent regulation of young FL-L1s by DNA methylation and MORC2.

Taken together, these data demonstrate that loss of DNA methylation activates transcription of young FL-L1s, which become targets of MORC2 (**Fig. 4G,H**). Simultaneous loss of DNMT1 and MORC2 therefore causes a massive accumulation in the levels of young FL-L1 transcripts, suggesting independent transcriptional and co/post-transcriptional L1 control by the two epigenetic pathways.

### HUSH-MORC2 and DNA methylation independently regulate ALRs and clustered protocadherins in hNPCs

Having established that DNA demethylation sensitizes L1 elements to HUSH-MORC2 restriction, we next considered whether this was a general effect at other classes of repetitive element. In subfamily-level analysis of RNA-seq data upon loss of MORC2 or TASOR individually, the alpha-satellite-like repeat (ALR) was by far the most upregulated class (**Fig. 5A**). We therefore considered the epigenetic status of ALRs. Although ultra-long (>100-kb) reads are required to uniquely map higher order arrays of ALRs at centromeres,^43^ the more scattered distribution of ALRs in pericentromeric regions allowed for unique mapping of thousands of our Nanopore reads (N50, 14-kb) that span relatively short ALR arrays and/or contain unique flanking sequences to aid mapping. We found that CpGs in pericentromeric regions containing ALRs were around 50% methylated, significantly below the whole genome average (**Fig. 5B,C**). Notably, ALRs were also markedly upregulated in hNPCs lacking DNMT1, illustrating that CpG methylation is nonetheless involved in controlling ALR transcription in hNPCs (**Fig. 5D**). DNA methylation status was unchanged over ALRs in cells depleted of MORC2 – suggesting that the pathways operate independently, as was observed at L1s (**Supp. Fig. 5A**). This conclusion was strengthened by the observation of a compounded effect on ALR transcript levels upon simultaneous depletion of MORC2 and DNMT1 (**Fig. 5D**). Together these results show that: (i) ALRs are hypomethylated and transcribed in hNPCs despite likely being embedded in a heterochromatin environment and (ii) HUSH-MORC2 controls ALR transcript levels at these regions. Since ALRs are markedly less methylated than the rest of the genome, and DNMT1 depletion further upregulates ALR transcription, our data would predict that robust DNA methylation of pericentromeric ALRs would silence these repeats and render them insensitive to loss of HUSH-MORC2. Whether particular sequence features (e.g. high AT-content) of ALRs underlie the sensitivity of these repeats to HUSH-MORC2, or change the dynamic between CpG methylation and repeat transcription at ALRs, requires further investigation.

**Figure 5.**
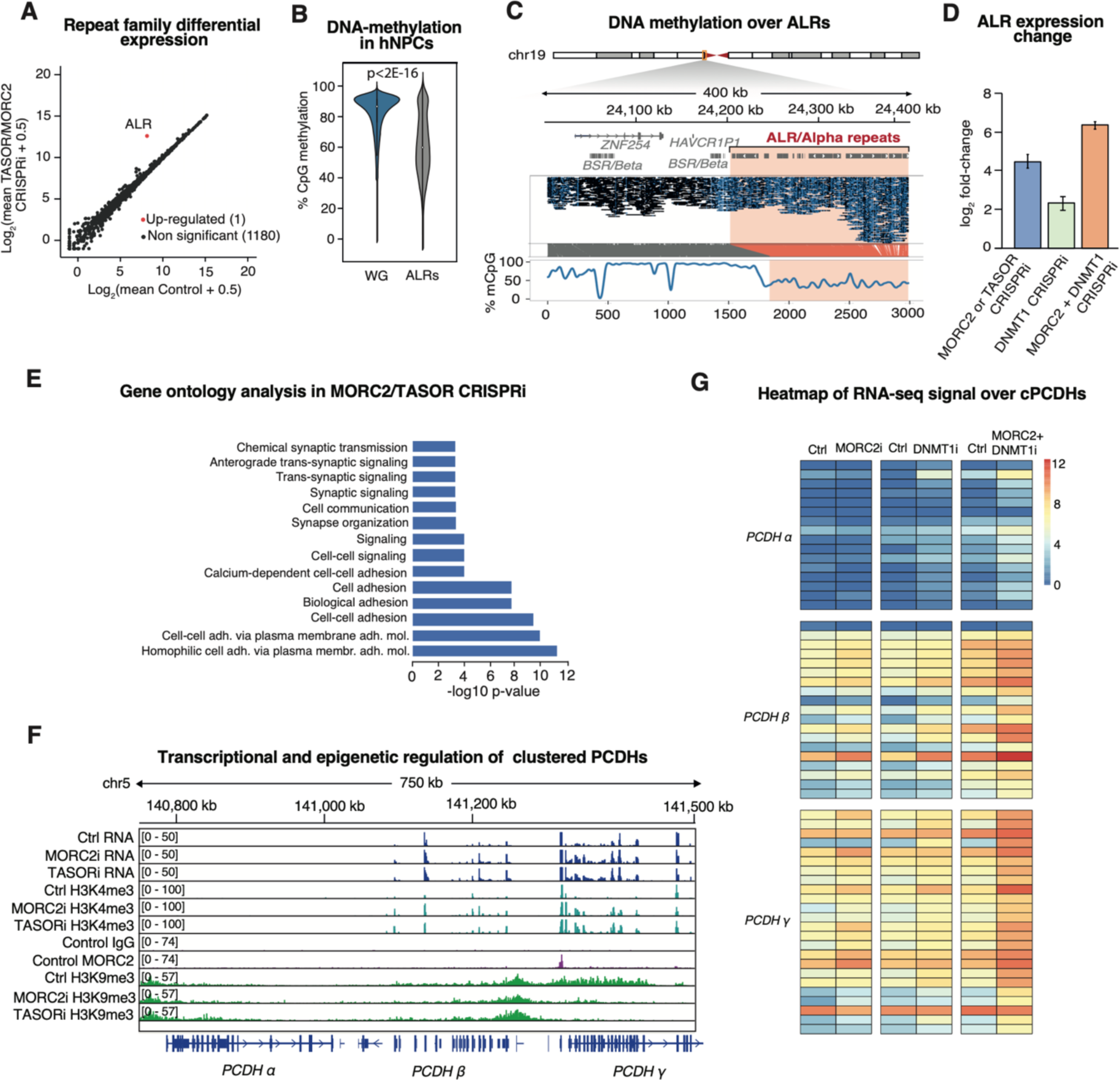
HUSH-MORC2 and DNA methylation independently control ALRs and protocadherins. **(A)** Mean plots of RNA-seq data in Control (n=4) and MORC2 and TASOR CRISPRi hNPCs (n=7) illustrating differential repeat expression based on TEtranscripts. ALR is the only subfamily with |log_2_ fold-change| >2 and padj <0.05. **(B)** Average CpG methylation in reads mapping uniquely to pericentromeric regions relative to the genome average (10-kb bins), illustrating relative ALR hypomethylation (Wilcoxon rank sum test with continuity correction). **(C)** Example of DNA methylation status in ALR-rich pericentromeric region on chr19. **(D)** Average log2-fold-change (LFC) of ALRs measured by RNA-seq in MORC2 or TASOR CRISPRi hNPCs (n=7, blue), DNMT1 CRISPRi (n=4, green) and MORC2/DNMT1 double CRISPRi (n=4, orange) versus controls (n=4 for each experiment). In each case the error bars represent +/- standard error of the mean calculated by DEseq2. **(E)** Gene ontology analysis of 373 differentially-expressed genes in MORC2 and TASOR CRISPRi hNPCs (no LFC cutoff, DEseq2 padj < 0.05). **(F)** Genome browser snapshot of the clustered protocadherin locus on chr5 illustrating transcriptional and chromatin regulation by HUSH-MORC2. **(G)** Heatmaps of clustered *PCDH* gene expression.

Lastly we considered the effect of HUSH-MORC2 on gene regulation in hNPCs. Of the 24 most upregulated genes in hNPCs lacking MORC2 or TASOR (LFC >2, **Supp. Fig. 5B**), 10 were in (ALR-rich) pericentromeric regions (**Supp. Fig. 5C**). Genes within 1MB of autosomal centromeres were significantly more strongly upregulated than those outside these regions (**Supp. Fig. 5D**). Gene ontology analysis of all significantly upregulated genes (n=373) revealed an enrichment of terms such as cell-cell adhesion and synapse organization (**Fig. 5E**), driven by protocadherin genes from the beta (*PCDHβ*) and gamma (*PCDHγ*) clusters on chromosome 5. This was accompanied by a gain of H3K4me3 over existing transcriptional start sites and the appearance of several new H3K4me3 peaks in both beta and gamma clusters (**Fig. 5F**). We observed near-total ablation of H3K9me3 over the *PCDHγ* cluster upon loss of MORC2 or TASOR but notably no change over the *PCDHβ* cluster, despite transcriptional and H3K4me3 changes over both clusters (**Fig. 5F**). MORC2 binding itself was identified at the locus, dominated by a large peak over *TAF7*, a single exon gene known to be HUSH-targeted^21^ that lies between the beta and gamma *PCDH* clusters (**Fig. 5F**). CUT&RUN data generated from bulk fetal forebrain and neurons from post-mortem adult cortical tissue (n=3) showed a large (∼750-kb) H3K9me3 domain over all three *PCDH* clusters (**Supp. Fig. 6A**). While we cannot prove this domain is HUSH-dependent in the human brain in vivo, we note that the complex is robustly expressed in both fetal and adult forebrain tissue according to in-house snRNA-seq data (**Supp. Fig. 6B, Fig. 1B, Supp. Fig. 1C**).^8, 44^

Since the transcriptional output of ALR and L1 repeats in hNPCs is independently controlled by DNA methylation and MORC2, we finally asked whether the same mechanistic hierarchy also applied at *PCDH*s. Loss of DNMT1 upregulated *PCDH* transcription and promoter H3K4me3 without affecting local H3K9me3 (**Supp. Fig. 6C**). MORC2 CRISPRi did not affect DNA methylation over the *PCDH* cluster (**Supp. Fig. 6D**). The double knockdown of MORC2 and DNMT1 caused greater transcriptional output from *PCDH*s than either depletion individually (**Fig. 5G**). Together these data suggest that *PCDH*s, like L1s and ALRs, are subject to a dual layer of epigenetic control in the developing human nervous system.

## DISCUSSION

As well as defending vertebrate genomes by silencing diverse exogenous genetic elements, the HUSH complex participates in the epigenetic control of endogenous repeats including L1 retrotransposons and repetitive genes.^7, 21, 22, 30, 45–47^ HUSH recruits the nuclear ATPase MORC2 that, together with H3K9me3-writer SETDB1, remodels chromatin at target loci.^7, 23, 25, 28, 48^ Recently, a series of studies have shown that HUSH impacts the transcriptional output of its targets by binding to nascent RNA and to nuclear complexes responsible for transcript degradation and termination.^26–28, 30^ However, a significant context to these findings is that they have been made in cells with hypomethylated genomes. For example, HUSH-MORC2 was originally identified as an L1 regulator in K562 cells,^7^ where the L1 promoter is essentially completely demethylated.^49^ Since DNA methylation is strongly correlated with transcriptional silencing of repeats,^17, 50^ and transcription is a critical determinant of HUSH-MORC2 targeting, whether and how these two pathways cooperate in the context of epigenetic repeat regulation is an important question that has remained largely unanswered.

In this study we systematically tested the relationship between HUSH-MORC2 and DNA methylation in controlling transcription of endogenous genomic repeats and repetitive genes in the human genome. To do this we took advantage of a somatic cellular model of early brain development where DNA methylation is robustly established but which, surprisingly, remains proliferative upon global genome demethylation by DNMT1 knockout.^36^ Through locus-specific analysis integrating Oxford Nanopore DNA methylation with bulk transcriptomic and histone methylation profiling, we identified that a handful of partially-demethylated L1 copies are weakly-transcribed and subject to restriction by HUSH-MORC2, while the vast majority are kept silent by DNA methylation. We therefore considered that genome demethylation should attract HUSH-MORC2 to activated L1 sequences. Indeed, upon loss of DNMT1 we could measure accumulation of MORC2 over activated young L1 integrants, and simultaneous CRISPRi-based depletion of DNMT1 and MORC2 led to massive accumulation of transcripts from these elements. We therefore demonstrate that DNA methylation controls the sensitivity of L1s to HUSH-MORC2 restriction and that these pathways operate independently to repress L1s.

In addition to L1 regulation, we identified the same mechanistic hierarchy between HUSH-MORC2 and DNA methylation in the transcriptional regulation of pericentromeric alpha-satellite-like repeats (ALRs). ALRs are hypomethylated, transcribed and targeted by HUSH-MORC2 at steady-state in hNPCs. Why other repeat transcripts (e.g. SVA retrotransposons and certain LTR elements, activated in cells lacking DNMT1) remain ignored by HUSH-MORC2 remains an open question. Details of ALR transcript structures and promoters remain largely uncharacterized, but two features of ALRs: an A-rich sequence and long transcriptional units, now seem to be characteristic of HUSH-sensitive elements^30^. More work is required to understand this specificity. While functional analyses of ALRs are sparse, transcription of these arrays is a conserved property that is thought to be important to centromere function.^51, 52^ An ever-improving range of long-read sequencing tools promises more functional studies of ALR and other tandem repeat transcription in health and disease, and HUSH will be a key player to consider here. In this regard it is interesting that despite HUSH being vertebrate-specific, it shares structural homology with the RITS complex^22^ that drives centromeric heterochromatin formation in yeast. MORC proteins are also found in basal eukaryotes and prokaryotes,^53^ suggesting that HUSH-like proteins with functions in silencing of repeats might form an ancient family with a conserved function in genome maintenance.

Regardless of its evolutionary origins, our observations have implications for the role of HUSH-MORC2 in hypomethylated contexts throughout human development. Predicted loss of function alleles in all members of the complex are strongly selected against according to aggregated analysis of more than 120,000 human exome sequences.^54^ It was recently hypothesized that HUSH-MORC2 might provide a compensatory mechanism during the wave of DNA demethylation that occurs after fertilization.^55^ In mouse embryonic stem cells (mESCs), DNA methylation is variable depending on culture conditions and – unlike human ESCs^56^ – can be removed via triple knockout of *Dnmt1, Dnmt3a* and *Dnmt3b*.^57^ Knockout of HUSH factor MPP8 caused cell cycle arrest in mESCs with low or absent DNA methylation, which was accompanied by the activation of murine L1 families.^58^ Thus, catastrophic repeat transcription (L1s or ALRs) and accompanying genome instability is a plausible explanation for the early developmental arrest phenotype in mice lacking TASOR^59^ or MORC2^60^ homologues. While the repetitive mouse genome (and L1 promoter structure) is divergent from that of humans and the consequences of DNA demethylation in mouse and human ESCs are different,^56^ our study provides direct evidence that HUSH-MORC2 protects the human genome from repeat activation in globally-hypomethylated cells or at specific hypomethylated loci. This protective mechanism in healthy tissue may be usefully subverted in cancer: indeed, in acute myeloid leukemia and solid tumor models – both characterized by aberrant DNA methylation – depletion of MPP8 or HUSH effector SETDB1 caused activation of tumor-suppressive transposons.^61, 62^ The therapeutic opportunity of HUSH-MORC2 inhibition in many cancers remains to be tested.

Finally, our observations of HUSH-MORC2 function in neural progenitors, complemented by epigenomic datasets of fetal and adult forebrain heterochromatin, have direct relevance in human neurobiology. Together with a recent study,^29^ we identified the HUSH-MORC2-H3K9me3 axis as a transcriptional regulator of clustered protocadherin genes. Stochastic promoter activation drives combinatorial *PCDH* expression in the nervous system, where each neuron displays a unique combination of PCDH molecules. How this beautiful genetic system is controlled by epigenetic mechanisms has been a fundamental question but is thought to involve a combination of DNA methylation and 3D structures influenced by SETDB1-dependent H3K9me3-heterochromatin.^63, 64^ We found that HUSH-MORC2 operates and does so independently of DNA methylation at this locus. This mechanism likely has clinical relevance: MORC2 ATPase domain mutations, which cause pronounced changes to the protein’s biochemical properties and HUSH activity at transgenes,^48^ cause neurodevelopmental disorders.^65, 66^ Since PCDHs drive the formation and complexity of neuronal networks and polymorphisms at the locus are themselves associated with a range of neurodevelopmental symptoms,^15^ (mis-)regulation of these genes by MORC2 mutants is an attractive explanation for the etiology of MORC2-disorders, though this remains to be tested. HUSH-MORC2 restriction of demethylated L1s is also likely relevant in the human brain. We recently reported widespread, cell-type-specific transcription of evolutionarily-young L1s in the developing and adult cortex,^8^ which may correlate with DNA methylation changes that occur in post-mitotic neurons.^67, 68^ How HUSH (and H3K9me3-heterochromatin more broadly) operates in regulation of L1s and other repeats during human ageing is an exciting open question.

## MATERIALS AND METHODS

### Antibodies

Rabbit anti-MORC2 (A300-149A, Bethyl; 1:1,000 dilution), rabbit anti-TASOR (Atlas HPA006735; 1:500), mouse anti-L1-ORF1p (Millipore MABC1152, 1:1,000) and HRP-conjugated anti-β-actin (Sigma A3854, 1:50,000) were used for Western blots. Rabbit anti-H3K4me3 (Active Motif 39159), rabbit anti-H3K9me3 (abcam 8898), rabbit anti-MORC2 (as above) and goat anti-rabbit IgG (abcam ab97047) were used for CUT&RUN assays (1:50-1:100 dilution). Rabbit anti-Nestin (Millipore AB5922, 1:100), mouse anti-beta-III-tubulin (Biolegend 801202, 1:500), chicken anti-GFP (abcam 13970, 1:100), mouse anti-5mC (Active Motif 39649, 1:250), Alexa647 anti-rabbit (Jackson Immuno Research 711-605-152, 1:500), Alexa647 anti-mouse (Jackson Immuno Research 115-605-003, 1:500) and Alexa488 anti-chicken (Jackson Immuno Research, 703-546-155, 1:500) were used for immunocytochemistry.

### Cell culture

The embryo-derived human neural progenitor cell line Sai2, which have the characteristics of neuroepithelial-like stem cells,^37^ was cultured according to standard protocol.^69^ Briefly, the cells were grown in DMEM/F12 (Thermo Fisher Scientific) supplemented with glutamine (Sigma), Penicillin/Streptomycin (1x, Gibco), N2 (1x, Thermo Fisher Scientific), B27 (0.05x, Thermo Fisher Scientific), EGF and FGF2 (both 10ng/ml, Thermo Fisher Scientific). Cells were cultured in flasks or multi-well plates (Nunc) pre-coated with poly L-ornithine (15μg/ml, Sigma) and laminin (2μg/ml, Sigma), fed daily with growth factors and passaged every 2-3 days using TrypLE Express enzyme (GIBCO) to detach cells and trypsin inhibitor (GIBCO) to quench the enzyme. 10 μM Rock inhibitor (Miltenyi) was added to the media following cell thawing, lentiviral transduction and FACS. Neural differentiation followed a detailed protocol for iPSC-derived hNPCs^70^. Cultures were tested routinely for Mycoplasma infection using Eurofins Mycoplasmacheck and returned a negative test.

### Human tissue

Human fetal forebrain tissue was obtained from material available following elective termination of pregnancy at the University Hospital in Malmö, Sweden, in accordance with the national ethical permit (Dnr 6.1.8-2887/2017). Post-mortem cortical tissue was obtained from the University Hospital in Lund, Sweden, in accordance with the national ethical permit (Dnr 2019-06582, 2020-02-12).

### Lentiviral vectors

Lentiviral vectors were produced according to a standard protocol.^71^ Briefly, low-passage 293T cells were grown to a confluency of ∼80% and co-transfected with three packaging plasmids (pMDL, psRev, and pMD2G) plus the relevant transfer plasmid using PEI. 48 h after transfection the virus-containing supernatant was collected, filtered, and centrifuged at 25,000xg for 1.5 h at 4 °C. The pellets were gently resuspended in PBS, aliquoted and stored at -80 °C until use. Titres were around 10^9^ TU/mL, determined by qPCR.

### CRISPR approaches

To inhibit the transcription of *MORC2, TASOR* and *DNMT1* individually, we followed a CRISPR interference approach described elsewhere.^72^ Single gRNA sequences were designed to bind just downstream of the relevant TSS and are listed below. Guides were inserted into a dCas9-KRAB-T2A-GFP lentiviral backbone containing both the gRNA under the U6 promoter and dCas9-KRAB-T2A-GFP under the Ubiquitin C promoter (pLV hU6-sgRNA hUbC-dCas9-KRAB-T2a-GFP, a gift from Charles Gersbach, Addgene plasmid #71237) using annealed oligos and the BsmBI cloning site. A control lentiviral vector expressing a gRNA sequence absent from the human genome (LacZ) was used as a control in all experiments. hNPCs were transduced with an MOI of 2.5-5 and GFP-positive cells FACS-isolated (FACSAria, BD sciences) on day 10 at 10°C. For CUT&RUN experiments, we avoided cell sorting so long as FACS analysis of the cell population indicated >90% GFP positives. For transcriptome analysis, sorted cells were pelleted at 400x*g* for 7 min, snap frozen on dry ice and stored at −80°C for RNA isolation.

For simultaneous depletion of DNMT1 and MORC2 we transduced with the same dCas9-KRAB-T2A-GFP vector expressing the most efficient MORC2 gRNA, together with the pLV.U6BsmBI.EFS-NS.H2b-mCh lentiviral backbone expressing the most efficient DNMT1 gRNA and an mCherry marker. Double positive cells were FACS sorted and pellets stored at -80 °C until RNA extraction.

For DNMT1 knockout experiments, LV.gRNA.CAS9-GFP vectors were used to target the catalytic domain of DNMT1 as described elsewhere.^36, 56^ Lentivirus was produced as above and used to transduce hNPCs at an MOI of 10–15. The same non-targeting control (LacZ) gRNA was used for the DNMT1 KO experiment, but in the relevant Cas9 backbone. As with CRISPRi experiments, GFP+ cells were isolated by FACS 10 days post-transduction.

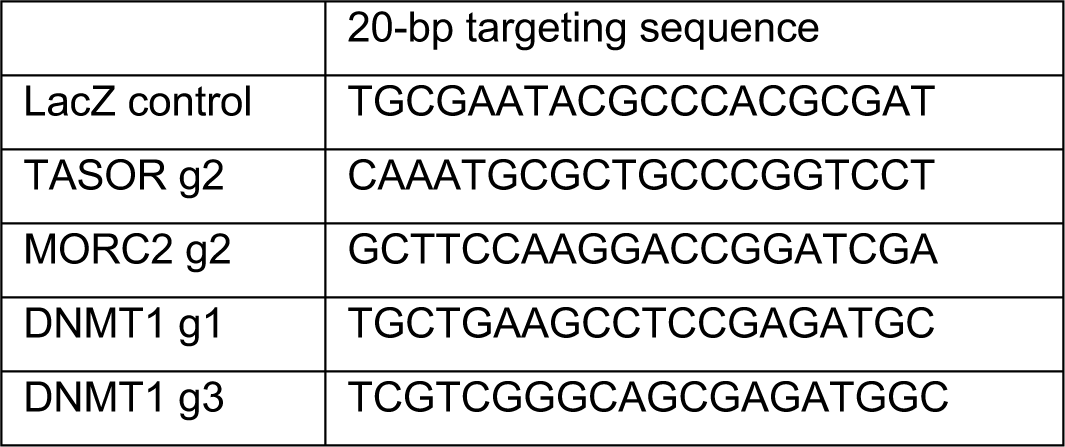

### Immunocytochemistry

Cells were gently rinsed in PBS three times and fixed in 4% formaldehyde for 15 min at room temperature. For blocking, cells were incubated for 1h with 5% Normal Donkey Serum (NDS) in TKBPS (KBPS with 0.25% Triton X-100) and then incubated (overnight, 4 °C) with the primary antibody (see above) or TKPBS + 5% NDS for negative controls. The following day cells were washed twice in TKPS then once in TKBPS/NDS, then incubated (2 h, room temperature) with the relevant secondary antibody (see above) and 5 min with DAPI (1:1000, Sigma D817) as a nuclear counterstain. For 5mC staining cells were pre-treated with 0.9% Triton in PBS (15 min) followed by 2 N HCl (15 min) then 10mM Tris pH 8 (10 min) prior to incubation with the primary antibody. Following washes in KPBS, cells were imaged using a fluorescence microscope (Leica) or an Operetta CLS High Content Analysis imager (PerkinElmer).

### RNA sequencing

Total RNA was isolated using the RNeasy Mini Kit (QIAGEN) with on-column DNAse treatment. Isolated RNA was used for RT-qPCR (see below) and/or RNA sequencing. Libraries for RNA sequencing were generated using Illumina TruSeq Stranded mRNA library prep kit (poly-A selection), optimized for long fragments by reducing fragmentation time, and sequenced on a NovaSeq6000 (paired end, 2×150 bp).

#### Quantification of genes

STAR RNA-seq aligner was employed to map the paired end reads to the human genome (GRCh38), using the annotations from gencode v38. Two parameters were modified from the default in STAR v2.6.0,^73^ namely increasing the number of allowed multimappers for each read (--outFilterMultimapNmax 100) and increasing the number of loci anchors (--winAnchorMultimapNmax 200). A sorted BAM was specified as the output which was further indexed using SAMtools v1.9.^74^ Finally, bamCoverage v2.4.3^75^ was used to create normalized tracks with a scaling factor of reads per kilobase million (RPKM). Reads from the BAM files were then quantified over the annotations from gencode v38 using subread’s featureCounts v1.6.3,^76^ specifying a parameter to force the strandedness of each feature to be taken into consideration (-s2).

#### Quantification of transposable elements

STAR RNA-seq v2.6.0 aligner was employed to map the paired end reads to the human genome (GRCh38), using the annotations from gencode v38. To obtain uniquely mapped reads, two of the default parameters were modified, where we allow each read to map to a single locus (--outFilterMultimapNmax 1) and allowing mismatch ratio of 0.03 (--outFilterMismatchNoverLmax 0.03). The resulting BAMs were indexed using SAMtools v1.9 and then used to create coverage tracks using bamCoverage v2.4.3 where an RPKM scaling factor was implemented. Finally, the uniquely mapped reads were quantified using featureCounts v1.6.3, specifying a parameter to force the strandedness of each feature to be taken into consideration (-s2). The multimapped reads were obtained identically to the BAMs in the gene quantification process and further quantified using TEtranscripts v2.0.3.^41^ To maintain consistency in our analyses, both the uniquely and multimapped reads were quantified over a curated GTF file provided by the creators of TEtranscripts.^41^

#### Differential expression analyses

Differential expression analyses for all bulk RNAseq data were performed with DESeq2 v1.38.3^77^ using the read count matrix from featureCounts as the input. Standard DESeq2 parameters were implemented where the counts were normalized by median of ratios. Visualization of the mean plots was achieved by obtaining the mean of normalized counts per condition, adding a pseudo count of 0.5 and log2 transforming the values.

### CUT&RUN

Three CUT&RUN protocols were used depending on the sample and target. A ‘standard’ protocol was used for profiling of histone modifications (H3K4me3, H3K9me3) in control or CRISPR-modified hNPC lines. A ‘nuclear’ protocol was used for profiling fetal and adult tissue samples. A ‘crosslinked’ protocol was used to profile MORC2 chromatin binding. *Standard CUT&RUN.* We followed the protocol described elsewhere^78^. Briefly, 250-500k cells were washed twice (20 mM HEPES pH 7.5, 150 mM NaCl, 0.5 mM spermidine, 1× Roche cOmplete protease inhibitors) and attached to 10 uL ConA-coated magnetic beads (Bangs Laboratories) that had been pre-activated in binding buffer (20 mM HEPES pH 7.9, 10 mM KCl, 1 mM CaCl2, 1 mM MnCl2). Bead-bound cells were resuspended in 50 mL buffer (20 mM HEPES pH 7.5, 0.15 M NaCl, 0.5 mM Spermidine, 1× Roche complete protease inhibitors, 0.05% w/v digitonin, 2 mM EDTA) containing primary antibody (see above) and incubated at 4 °C overnight with gentle rotation. Beads were washed thoroughly with digitonin buffer (20 mM HEPES pH 7.5, 150 mM NaCl, 0.5 mM Spermidine, 1× Roche cOmplete protease inhibitors, 0.05% digitonin). After the final wash, pA-MNase (a generous gift from Steve Henikoff) was added in digitonin buffer and incubated with the cells at 4 °C for 1 h. Bead-bound cells were washed twice, resuspended in 100 uL digitonin buffer, and chilled to 0-2 °C. Genome cleavage was stimulated by addition of 2 mM CaCl2 at 0 °C for 30 min. The reaction was quenched by addition of 100 mL 2× stop buffer (0.35 M NaCl, 20 mM EDTA, 4mM EGTA, 0.05% digitonin, 50 ng/mL glycogen, 50 ng/mL RNase A, 10 fg/mL yeast spike-in DNA (a generous gift from Steve Henikoff)) and vortexing. After 10 min incubation at 37 °C to release genomic fragments, cells and beads were pelleted by centrifugation (16,000 g, 5 min, 4 °C) and fragments from the supernatant were purified by PCR clean-up spin column (Macherey-Nagel).

#### Crosslinked CUT&RUN

The procedure was the same as above except for the following changes: 900K-1M cells were used and crosslinked for 1 min using 1% formaldehyde (16% solution, Thermo Scientific) diluted in media. Cells were washed three times (20 mM HEPES pH 7.5, 150 mM NaCl, 0.5 mM spermidine, 1× Roche complete protease inhibitors, 1% Triton X-100, 0.05% SDS) with a slightly increased centrifugation speed to avoid pellet loss. After attachment to ConA beads, the mixture was resuspended in buffer (20 mM HEPES pH 7.5, 150 mM NaCl, 0.5 mM spermidine, 1× Roche complete protease inhibitors, 1% Triton X-100, 0.05% SDS, 0.05% digitonin, 2 mM EDTA) containing primary antibody. Following 4 °C overnight incubation with primary antibodies, beads were washed thoroughly with digitonin buffer (20 mM HEPES pH 7.5, 150 mM NaCl, 0.5 mM Spermidine, 1× Roche complete protease inhibitors, 1% Triton X-100, 0.05% SDS, 0.05% digitonin). After incubation at 37 °C to release genomic fragments, crosslinking was reversed by adding 0.09% SDS and 0.22 mg/ml proteinase K (Thermo Scientific) and incubation overnight at 55 °C. The following day, DNA was purified by spin column as above.

#### Nuclear CUT&RUN

Isolation of nuclei from embryonic and adult human brain tissue was performed according to an established protocol.^79^ In brief, tissue was dissociated in ice-cold lysis buffer (0.32 M sucrose, 5 mM CaCl2, 3 mM MgOAc2, 0.1 mM Na2EDTA, 10 mM Tris-HCl, pH 8.0, 1 mM DTT) using a 1-mL Dounce homogenizer (Wheaton). The homogenate was carefully layered on a sucrose cushion (1.8 M sucrose, 3 mM MgOAc2, 10 mM Tris-HCl, pH 8.0, and 1 mM DTT) before centrifugation (30,000×*g*, 2 h 15 min). Pelleted nuclei were softened for 10 min in 100 mL of nuclear storage buffer (15% sucrose, 10 mM Tris-HCl, pH 7.2, 70 mM KCl, and 2 mM MgCl2) then resuspended in 300 mL of dilution buffer (10 mM Tris-HCl, pH 7.2, 70 mM KCl, and 2 mM MgCl2) and run through a cell strainer (70 mm). Cells were run through the FACS (FACS Aria, BD Biosciences) at 4°C at a low flow rate using a 100 mm nozzle (reanalysis showed >99% purity). For isolation of NeuN+ nuclei (adult post-mortem samples only), nuclei were incubated with AlexaFluor 488 anti-NeuN (abcam 190195, 1:500) for 30 min on ice. 300k AlexaFluor488-positive nuclei were sorted per tissue piece. The sorted nuclei were pelleted at 1,300 × g for 15 min and resuspended in 1 mL of ice-cold nuclear wash buffer (20 mM HEPES, 150 mM NaCl, 0.5 mM spermidine, 1× cOmplete protease inhibitors, 0.1% BSA) and 10 µL per antibody treatment of ConA-coated magnetic beads (Epicypher) added with gentle vortexing. All buffers contained 0.1% BSA and tips were pre-treated with 0.1% BSA.

#### CUT&RUN analysis

Illumina sequencing libraries were prepared using the Hyperprep kit (KAPA) with unique dual indexed adapters (KAPA), pooled and sequenced on a Nextseq500 instrument (Illumina). Paired-end reads (2×75) were aligned to the human genome (hg38) using bowtie2 (–local –very-sensitive-local –no-mixed –no-discordant –phred33 -I 10 -X 700) and then converted converted to bam files using samtools. ^74, 80^ To extract the uniquely aligned reads, BAM files were filtered setting a threshold for mapping quality of 10 (MAPQ10). Coverage bigwig tracks were created using bamCoverage (deepTools)^75^ and normalized with a rounds per kilobase million (RPKM) scaling factor. The coverage tracks displayed in all figures were visualized by the genome browser IGV. Peaks were called using HOMER findPeaks v4.10,^81^ searching for peaks with variable lengths using the histone command (-style histone). In order to capture both baseline and knockdown specific peaks, we used the command for both conditions (control and CRISPRi) and filtered out all non-canonical chromosomes. H3K9me3 peaks were also filtered according to length, where all peaks shorter than 1 kb were excluded from the analysis. Finally, to make a comprehensive peak list, we concatenated and merged the baseline and knockdown specific peaks. Tag directories were created with HOMER’s makeTagDirectory v4.10 using the standard parameters. Finally, we obtained the differential enrichment over the peaks discovered by SEACR using HOMER’s getDifferentialPeaks setting a fold change threshold of 3 (-F 3) and keeping the default settings for the rest of the parameters. Heatmap matrices were computed using DeepTools’ computeMatrix v2.5.4 and later visualized with plotHeatmap 2.5.4 from the same deepTools package.^75^

### RT-qPCR

cDNA was generated by reverse transcription of 500ng RNA with random hexamer primers and Superscript III (Invitrogen) and analysed by RT-qPCR with SYBR Green I master (Roche) on a LightCycler 480 instrument (Roche). Data are represented with the ΔΔCt method normalized to housekeeping genes *ACTB* and *HPRT*. Primer sequences are listed below.

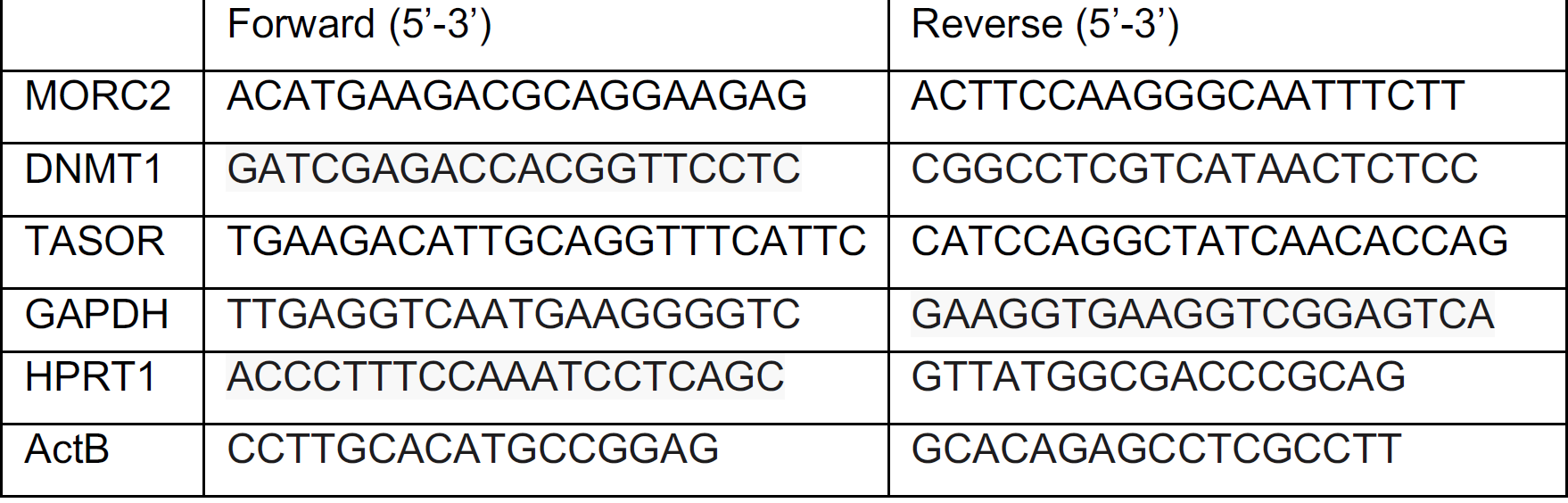

### Western blot

Cells were lysed in RIPA buffer (Sigma-Aldrich) containing complete protease inhibitor cocktail on ice for 30 min and then pelleted (20 min, 17,000x*g*, 4 °C). Supernatants were collected and mixed with Novex LDS 4× loading dye containing reducing agent (Thermo) boiled at 95 °C for 5 min before being separated on a 4–12% Tris-glycine SDS-PAGE gel (200 V, 45 min). Proteins were transferred from the gel to a PVDF membrane using the Transblot-Turbo Transfer system (BioRad). The membrane was then washed twice (15 min) in Tris-buffered saline with 0.1% Tween (TBST) and blocked for 1 h in TBST with 5% skimmed milk (MTBST) before incubation (overnight, 4 °C) with the primary antibody diluted in MTBST. The following day the membrane was washed twice in TBST (15 min) and incubated (1 h, room temperature) with HRP-conjugated anti-mouse or anti-rabbit secondary antibody (Santa Cruz Biotechnology, 1:5,000) diluted in MTBST. After washing the membrane twice in TBST and once in TBS, protein was detected by chemiluminescence using ECL Select reagents (Cytiva) as per manufacturer’s instructions and imaged on a Chemi-Doc system (BioRad). The membrane was stripped using the Restore PLUS Wester Blot Stripping Buffer (Thermo) as per instructions, re-blocked for 1 h in MTBST after which the procedure for the β-actin staining was performed as above.

### Oxford Nanopore sequencing

High molecular weight DNA was extracted from frozen pellets (1 million cells) using the Nanobind HMW DNA Extraction kit (PacBio) following the manufacturer’s instructions. The final product was eluted in 100 μL. DNA concentration and quality were measured using Nanodrop and Qubit from the top, middle, and bottom of each tube, and by agarose gel electrophoresis. Whole genome sequencing was done using SQK-LSK109 Ligation Sequencing kit (Oxford Nanopore Technologies) and FLO-PRO002 PromethION Flow Cell R9 Version and was sequenced on a PromethION (Oxford Nanopore Technologies) at SciLifeLab. Long read DNA sequencing data were aligned to the human genome (hg38) using minimap2 (-a -x map-ont)^82^ and 5mC base modifications were detected using the nanopolish tool.^83^ We then used methylartist to create methylation databases (methylartist db-nanopolish -m nanopolish.tsv.gz -d nanopolish.db), and further employed it for all visualizations.^84^

## DATA AND CODE AVAILABILITY

There are no restrictions on data availability. The RNA and DNA sequencing data presented in this study have been deposited at GEOs:

-GSEXXXXX: bulk RNA-seq, CUT&RUN and ONT whole genome sequencing of hNPC CRISPRi and control cells.

-GSE224747: 3’ single nuclei RNAseq, CUT&RUN and bulk RNAseq of fetal tissue samples.

-GSE209552: 3’ single nuclei RNAseq of adult samples.

-GSE211871: Adult post-mortem NeuN+ CUT&RUN sequencing.

## ACKNOWLEDGMENTS

We thank M. Persson-Vejgården, A. Hammarberg and E. Monni for their technical assistance, S. Henikoff for the gift of the pA-MNase used in CUT&RUN experiments, A. Falk for providing hNPC lines, and D. O’Carroll and D. Prigozhin for comments on the manuscript. This work was supported by grants from the Swedish Research Council (VR, 2018-02694 to J.J., 2020-01660 to Z.K. and 2021-03494 to C.H.D.) the Swedish Brain Foundation (Hjärnfonden, FO2019-0098 to J.J. and FO2022-0079 to Z.K), the Swedish Society for Medical Research (SSMF, S19-0100 to C.H.D.), the Crafoord Foundation, Hedlund Foundation, Jeansson Foundation (project grants to C.H.D.) and the Swedish Government Initiative for Strategic Research Areas (MultiPark & StemTherapy).

## AUTHOR CONTRIBUTIONS

N.P., J.J. and C.H.D. conceived and designed the study. V.H., O.E.K., G.C., F.D., S.K., J.M., D.A., J.G.J and C.H.D. performed experimental work. N.P., S.K., P.G. and R.G. performed bioinformatic analyses. M.E.J., A.A., E.E. and Z.K. contributed essential materials or expertise. C.H.D. wrote the manuscript with assistance from N.P. and V.H., who prepared the figures. J.J. and C.H.D. supervised the study and acquired funding. All authors reviewed the final version.

## SUPPLEMENTARY FIGURES

**Supplementary Figure 1.**
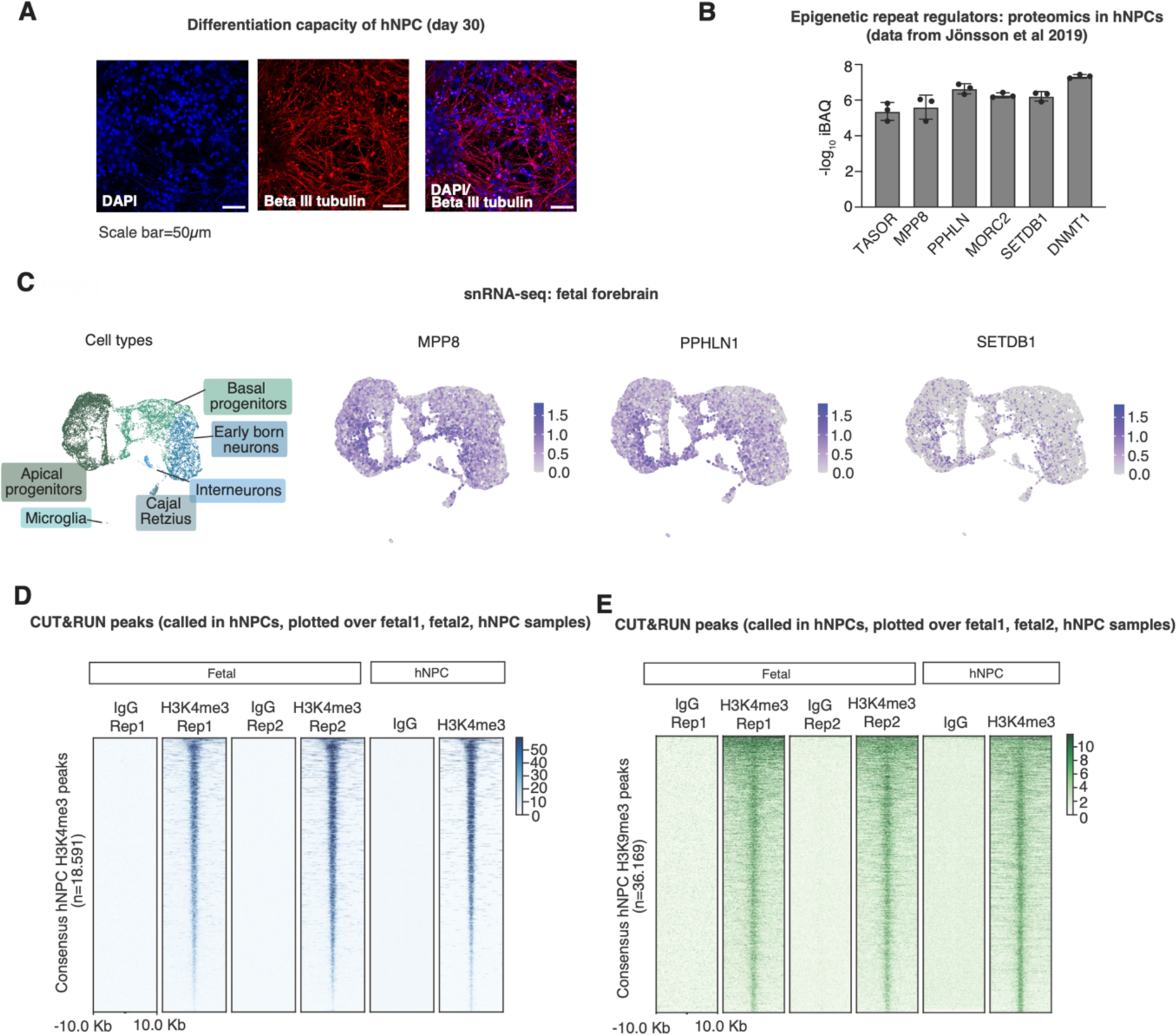
**(A)** Immunostaining of day 30 neurons differentiated from the hNPC line used in this study illustrating expression of neuronal marker beta-III-tubulin. (**B)** Detection of epigenetic repeat regulators in published hNPC proteomic analysis.^36^ **(C)** Expression of MPP8, PPHLN1 and SETDB1 in the developing human forebrain according to snRNA-seq analysis (n=6).^8^ **(D, E**) Heatmaps illustrating CUT&RUN signal enrichment of H3K4me3, H3K9me3 and a non-targeting IgG control in fetal forebrain samples and hNPCs plotted over consensus peaks called in hNPC samples. Displayed are the genomic regions spanning +/- 10kb from the peak centre.

**Supplementary Figure 2.**
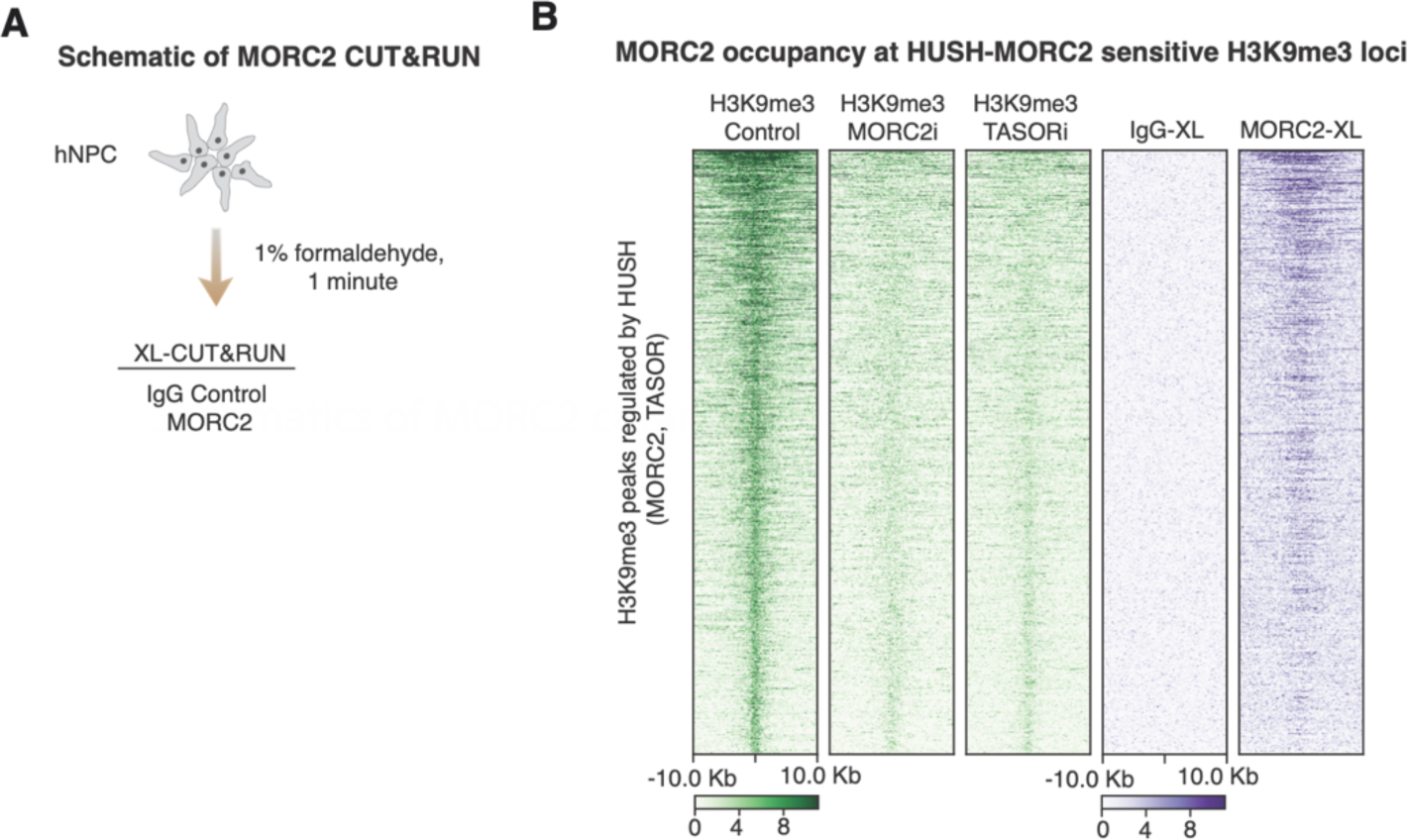
**(A)** Schematic of MORC2 CUT&RUN experiments in hNPCs using a crosslinking protocol. **(B)** Presence of MORC2 over TASOR and MORC2-controlled H3K9me3 peaks in hNPCs. Displayed are the genomic regions spanning +/- 10kb from the peak centre.

**Supplementary Figure 3.**
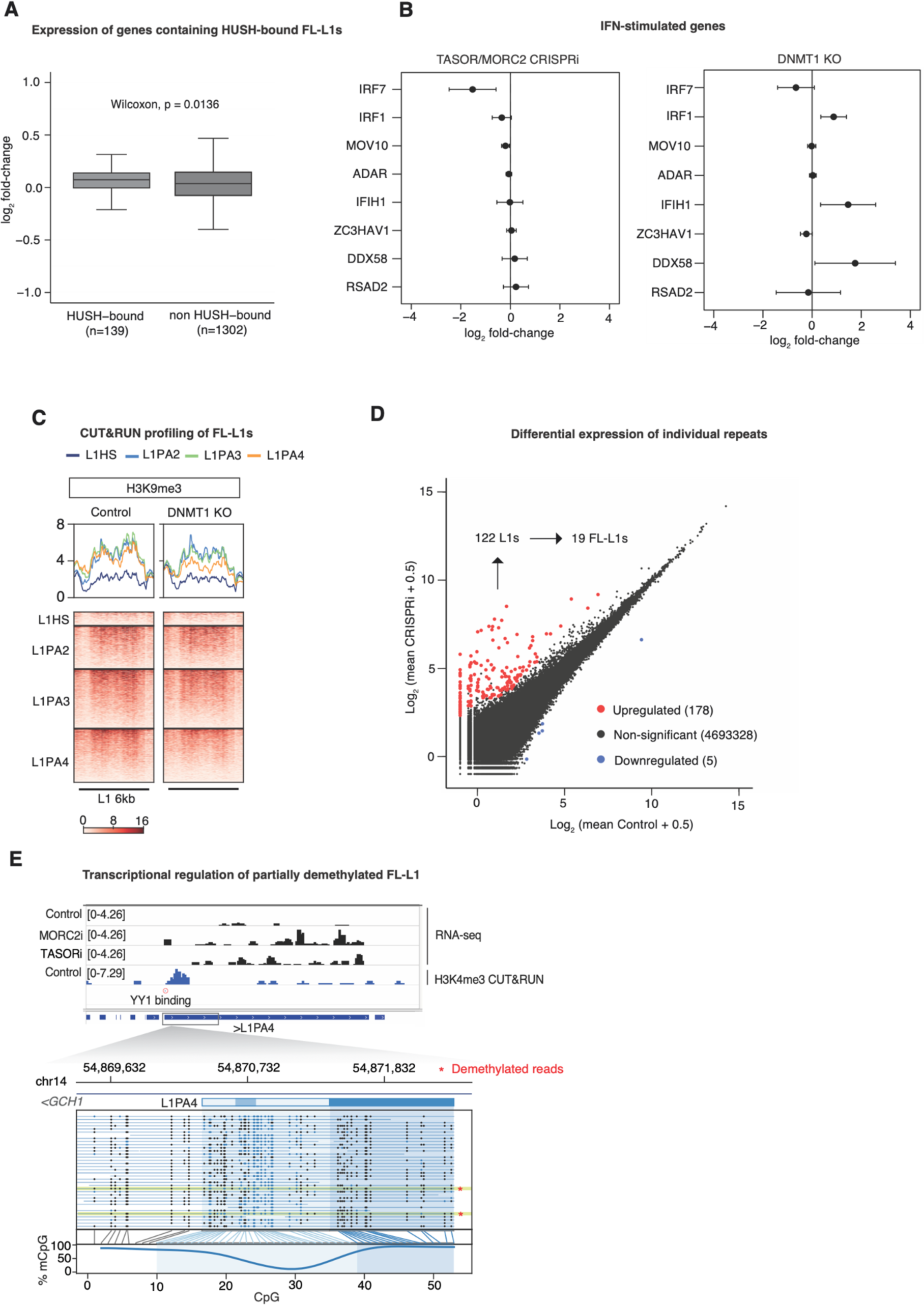
**(A)** Comparison of genes expression changes upon MORC2 and TASOR CRISPRi between genes containing FL-L1s marked by TASOR and MORC2-dependent H3K9me3 and genes containing non-targeted FL-L1s. The central bands denote medians. Boxes represent the interquartile range (IQR). Whiskers extend 1.5x IQR beyond the box. Statistical test: Wilcoxon rank sum and signed rank. **(B)** Differential expression analysis of selected interferon (IFN) stimulated genes in the TASOR and MORC2 CRISPRi treatments (n=7) versus control hNPCs (n=4) and in DNMT1-KO hNPCs (n=3) versus control (n=3). The DNMT1-KO RNA-seq data was taken from Jönsson et al.^36^ **(C)** CUT&RUN H3K9me3 signal plotted over evolutionarily-young FL-L1s sorted by evolutionary age in control and DNMT1-KO hNPCs. The experiment was repeated twice with similar results. Only uniquely-mapped reads were retained. **(D)** Mean plot illustrating differential expression analysis of individual repeats from RepeatMasker in MORC2 and TASOR CRISPRi treatments (n=7) versus control hNPCs (n=4). Elements with |log_2_ fold-change| >2 and padj <0.05 are highlighted. The number of upregulated L1s and FL-L1s are given. **(E)** A third example of a FL-L1 weakly upregulated by MORC2 and TASOR CRISPRi showing promoter H3K4me3 in control cells and an intact YY1 binding site (upper). Nanopore sequencing reads suggest incomplete DNA methylation over the promoter in control hNPCs (lower). See also Fig. 3.

**Supplementary Figure 4.**
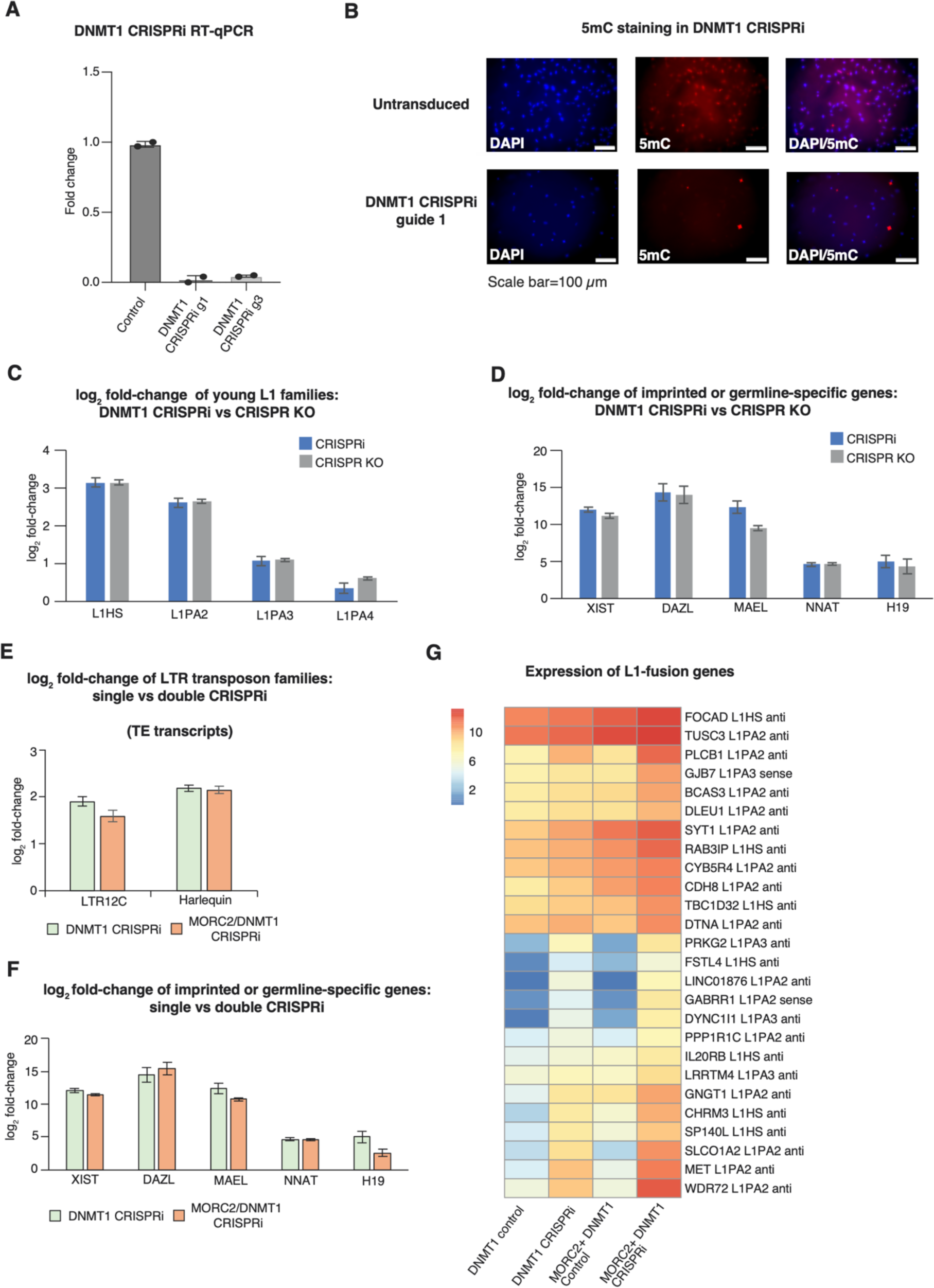
**(A)** Reverse transcription quantitative PCR (RT-qPCR) of DNMT1 transcript levels in DNMT1 CRISPRi cells, relative to a non-targeting control, 10 days post-transduction. **(B)** 5mC immunostaining in control and DNMT1 CRISPRi cells. **(C)** Average TEtranscripts log2-fold-change (LFC) of young L1 subfamilies in DNMT1 CRISPRi (n=4) versus control hNPCs (n=4) and DNMT1-KO (n=3) versus control hNPCs (n=3). DNMT1-KO data taken from Jönsson et al.^36^ **(D)** Average LFC of imprinted and germline-specific genes in DNMT1 CRISPRi (n=4) versus control hNPCs (n=4) and DNMT1-KO (n=3) versus control hNPCs (n=3). DNMT1-KO data taken from Jönsson et al.^36^ **(E)** Average TEtranscripts LFC of selected DNMT1-controlled LTR families measured by RNA-seq in DNMT1 CRISPRi (n=4, green) and MORC2/DNMT1 double CRISPRi (n=4, orange) versus controls (n=4 for each experiment). **(F)** Average LFC of imprinted and germline-specific genes in DNMT1 CRISPRi (n=4, green) and MORC2/DNMT1 double CRISPRi (n=4, orange) versus controls (n=4 for each experiment). In each case the error bars represent +/- standard error of the mean calculated by DEseq2. **(G)** Heatmap of normalized read counts of L1-fusion genes^36^ measured by RNA-seq in DNMT1 CRISPRi (n=4) and MORC2/DNMT1 double CRISPRi (n=4) versus controls (n=4 for each experiment). Given next to each row are the subfamily to which the L1 belongs and whether it occupies the same (sense) or opposite (anti) strand relative to the gene.

**Supplementary Figure 5.**
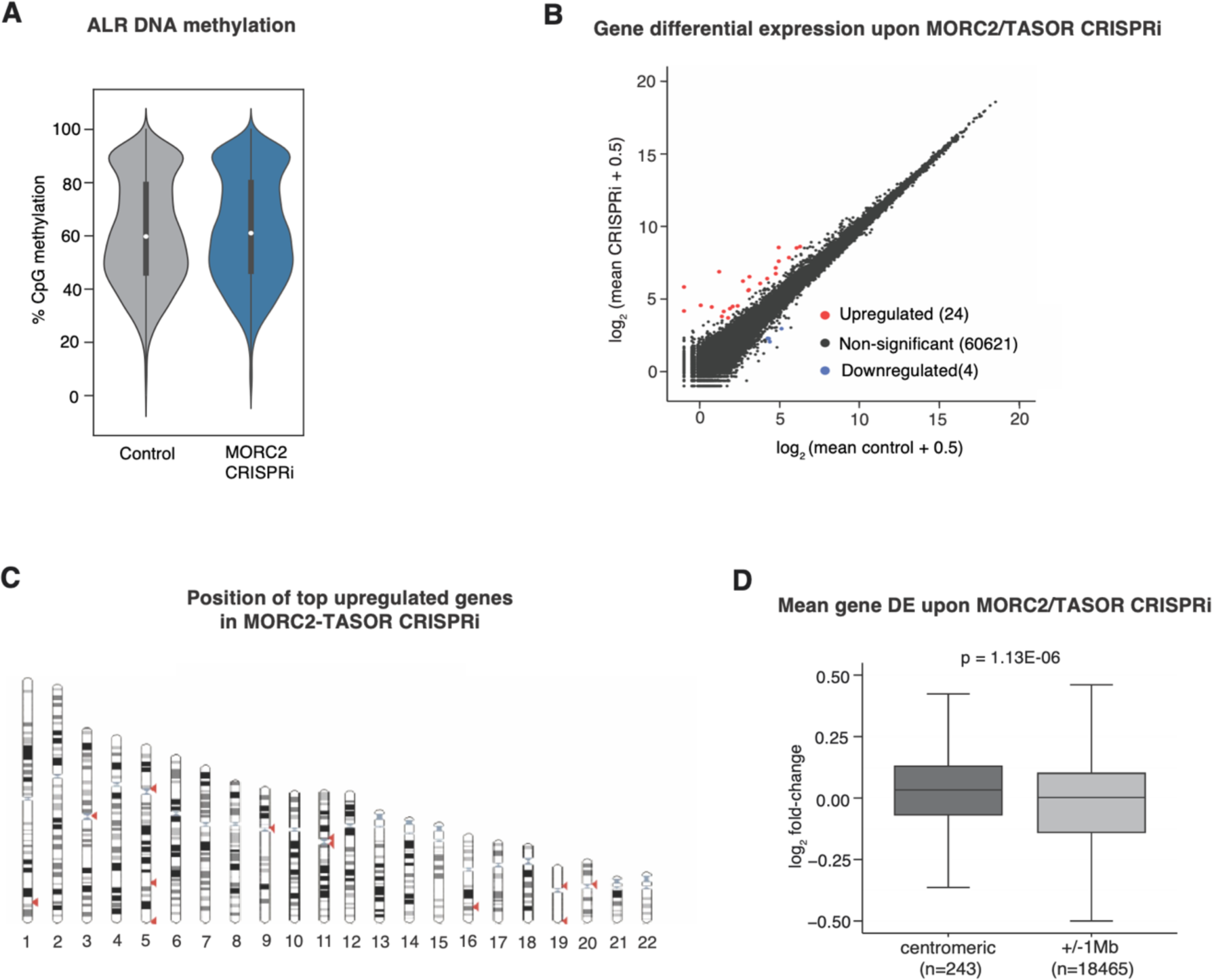
**(A)** Average CpG methylation in reads mapping uniquely to pericentromeric regions in whole-genome Nanopore sequencing data from control (n=1) and MORC2 CRISPRi hNPCs (n=1). **(B)** Differential gene expression in MORC2 and TASOR CRISPRi (n=7) versus control hNPCs (n=4). Elements with |log_2_ fold-change| >2 and padj <0.05 are highlighted. **(C)** Chromosome positions of top upregulated genes in MORC2 and TASOR CRISPRi hNPCs. **(D)** Comparison of genes expression changes upon MORC2 and TASOR CRISPRi between genes within 1 Mb of an autosome centromere (left) and those outside (right). The central bands denote medians. Boxes represent the interquartile range (IQR). Whiskers extend 1.5x IQR beyond the box. Statistical test: Wilcoxon rank sum with signed rank.

**Supplementary Figure 6.**
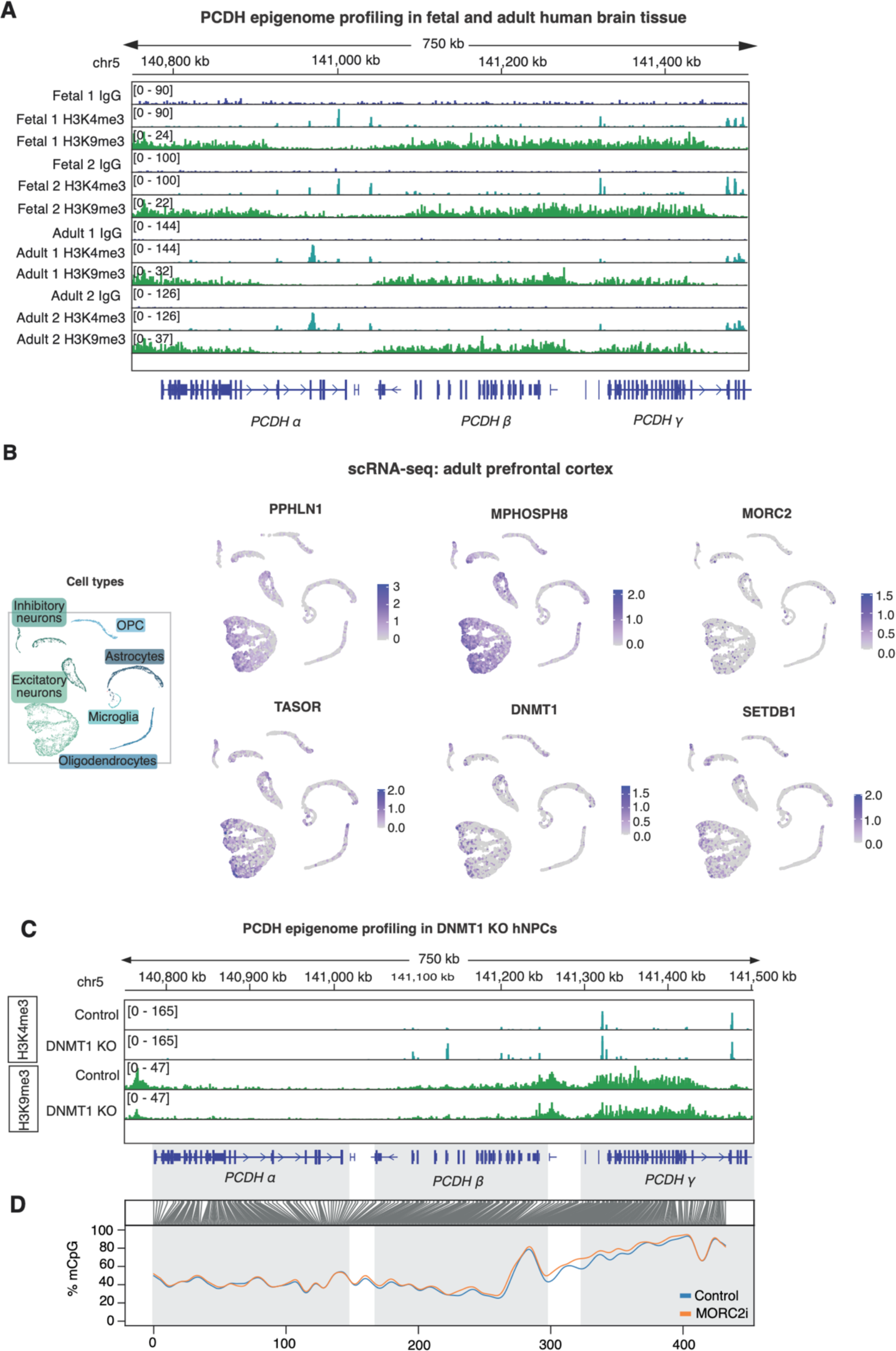
**(A)** Genome browser snapshot illustrating epigenomic profiling of clustered protocadherins (*PCDH*) in fetal and adult post-mortem human brain tissue. For adult tissue, NeuN+ nuclei were sorted prior to CUT&RUN. **(B)** Expression of epigenetic repeat regulators in snRNA-seq analysis of post-mortem human cortical tissue (n=5).^44^ **(C)** H3K4me3 and H3K9me3 profiles in control and DNMT1-KO hNPCs over clustered *PCDH* genes. Experiments were repeated twice with similar results. **(D)** DNA methylation profile of control (n=1) and MORC2 CRISPRi hNPCs (n=1) according to whole-genome Oxford Nanopore Sequencing analysis.

